# An intrinsically disordered region of Drosha selectively promotes miRNA biogenesis, independent of tissue-specific Microprocessor condensates

**DOI:** 10.1101/2025.04.10.648254

**Authors:** Bing Yang, Brian Galletta, Rima Sakhawala, Nasser Rusan, Katherine McJunkin

## Abstract

Precise control of miRNA biogenesis is of extreme importance, since mis-regulation of miRNAs underlies or exacerbates many disease states. The Microprocessor complex, composed of DROSHA and DGCR8, carries out the first cleavage step in canonical miRNA biogenesis. Despite recent advances in understanding the molecular mechanism of Microprocessor, the N-terminal region of DROSHA is less characterized due its high intrinsic disorder. Here we demonstrate that Microprocessor forms condensates with properties consistent with liquid-liquid phase separation (LLPS) in select tissues in *C. elegans*. While DRSH-1/Drosha recruitment to granules is only partially dependent on its intrinsically disordered regions (IDRs), one of these N-terminal IDRs is crucial for biogenesis of a subset of miRNAs and normal development. A cis region of IDR-dependent miRNAs confers IDR-dependence to another miRNA, suggesting that the IDR recognizes sequences or structures in the miRNA primary transcript. Future studies will further elucidate the specificity of this interaction and the putative role of Microprocessor condensates.

## Introduction

microRNAs (miRNAs) are a class of small non-coding RNAs around 22-23 nucleotides in length (Bartel, 2018). Their biogenesis starts with transcription of primary miRNA transcripts (Cai et al., 2004; H. Kim et al., 2024; Lee et al., 2004) which are then recognized and cleaved by the Microprocessor complex composed of DROSHA and DGCR8 (also known as PASH-1 in *C. elegans*) resulting in precursor miRNAs with sizes from ∼60 to 80nt (Denli et al., 2004; Gregory et al., 2004; Han et al., 2004, 2006; H. Kim et al., 2024; Landthaler et al., 2004; Lee et al., 2003; Zeng et al., 2005). The precursor miRNA is then exported from the nucleus to the cytoplasm by exportins (Bohnsack et al., 2004; Büssing et al., 2010; Lund et al., 2004; Yi et al., 2003; Zeng & Cullen, 2004). In the cytoplasm, endonuclease DICER recognizes and cleaves the precursor into a mature miRNA duplex (Bernstein et al., 2001; Grishok et al., 2001; Hutvágner et al., 2001; Ketting et al., 2001; Knight & Bass, 2001), which is then loaded into an Argonaute protein to form the miRNA induced silencing complex (miRISC) and target downstream mRNA molecules for gene silencing (Gregory et al., 2005; Grishok et al., 2001; Hammond et al., 2001; Mourelatos et al., 2002; Tabara et al., 1999).

Primary miRNA (pri-miRNA) recognition and cleavage is orchestrated by the Microprocessor complex (Denli et al., 2004; Gregory et al., 2004; Han et al., 2004, 2006; H. Kim et al., 2024; Landthaler et al., 2004; Lee et al., 2003; Zeng et al., 2005). Structural and *in vitro* biochemical studies have intensively characterized the interaction interfaces between the Microprocessor complex and the pri-miRNA molecule; this work has revealed several sequence elements that define substrate optimality of pri-miRNAs, such as the UG motif near the basal junction (Auyeung et al., 2013; Fang & Bartel, 2015; Garg et al., 2024; Jin et al., 2020; B. Kim et al., 2017; Kwon et al., 2016; Li et al., 2020; T. A. Nguyen et al., 2015; Partin et al., 2020). However, most of these studies were carried out using truncated DROSHA without the N-terminus due to aberrant aggregation of the full-length protein (T. A. Nguyen et al., 2015). Therefore, in contrast to the intensive characterization of the enzymatic domains, the function of the N-terminus of DROSHA is poorly understood. The N-terminal region of DROSHA is the least conserved region of the protein (B. Kim et al., 2017). In vertebrates, the N-terminus is highly enriched in prolines, arginines, and serines, and enriched in overall disorderedness (Son et al., 2023). Global miRNA deficiency was observed upon endogenous deletion of the DROSHA N-terminus (A.A. 1-396) through CRISPR in cell lines (Prabhakar et al., 2023). Another recent report identified a major proteolytic DROSHA isoform that lacks the N-terminal proline-rich disordered domain (PRD – A.A. 1∼198) and described the function of the PRD in mediating pri-miRNA processing of a subset of miRNAs encoded in introns (Son et al., 2023).

Over the last decade, liquid-liquid phase separation (LLPS) has been linked to numerous biological phenomena, and LLPS is thought to be the driving force behind the formation of a diverse array of membrane-less organelles, such as stress granules, P-bodies, paraspeckles, and P granules (Banani et al., 2017; Brangwynne et al., 2009). Scaffold molecules initiate the formation of the scaffold matrix through multivalent interactions and then recruit client molecules to further mature these condensates (Banani et al., 2016). Intrinsically disordered regions (IDR) of proteins are hypothesized to be one of the main providers of multivalency (Mitrea & Kriwacki, 2016). Liquid-like biomolecular condensates have been proposed to support diverse functions such as compartmentalization and concentration of biochemical reactions or storage of macromolecules (Putnam et al., 2023). A key question in the field is how frequently observable biomolecular phase-separated droplets represent “incidental condensates” that do not add functionality beyond that provided by the soluble population of proteins outside of the condensates (Putnam et al., 2023).

In this study, we describe nuclear condensate formation of the Microprocessor complex in select *C. elegans* tissues. FRAP experiments suggest that the Microprocessor foci are dynamic, consistent with formation through LLPS. Based on co-localization analysis, we conclude that these DRSH-1 condensates are distinct from known nuclear granules. We then investigate the function of the predicted IDRs in regulating DRSH-1 condensate formation and miRNA biogenesis. We observe that an IDR is required for the biogenesis of a subset of miRNAs and for normal development. However, the roles of the Drosha IDR in miRNA biogenesis and animal development appear to be independent of the condensate formation.

## Results

### Drosha forms distinct nuclear condensates in *C. elegans*

To assess the distribution of the Microprocessor in *C. elegans*, we fused GFP to the DRSH-1 C-terminus or mCherry to the PASH-1 C-terminus by knock-in at the respective endogenous locus using CRISPR/Cas9 (Figure 1A). To ensure the normal functionality of the fluorescently tagged proteins, we performed co-immunoprecipitation of DRSH-1::GFP and PASH-1::mScarlet and observed a strong interaction, as expected of a functional Microprocessor complex (Figure S1A). To further ensure that the tagged Microprocessor exhibits wild type function at the molecular level, we prepared small RNA libraries from these fluorescently tagged strains, including *drsh-1::gfp, pash-1::mScarlet* and *drsh-1::gfp pash-1::mScarlet*, and sequencing revealed no significant changes of miRNA levels when compared to wild type animals (Figure S1B-D, Table S16-18). Thus, these endogenously tagged Microprocessor components function in a wild type manner in miRNA biogenesis.

**Figure 1.**
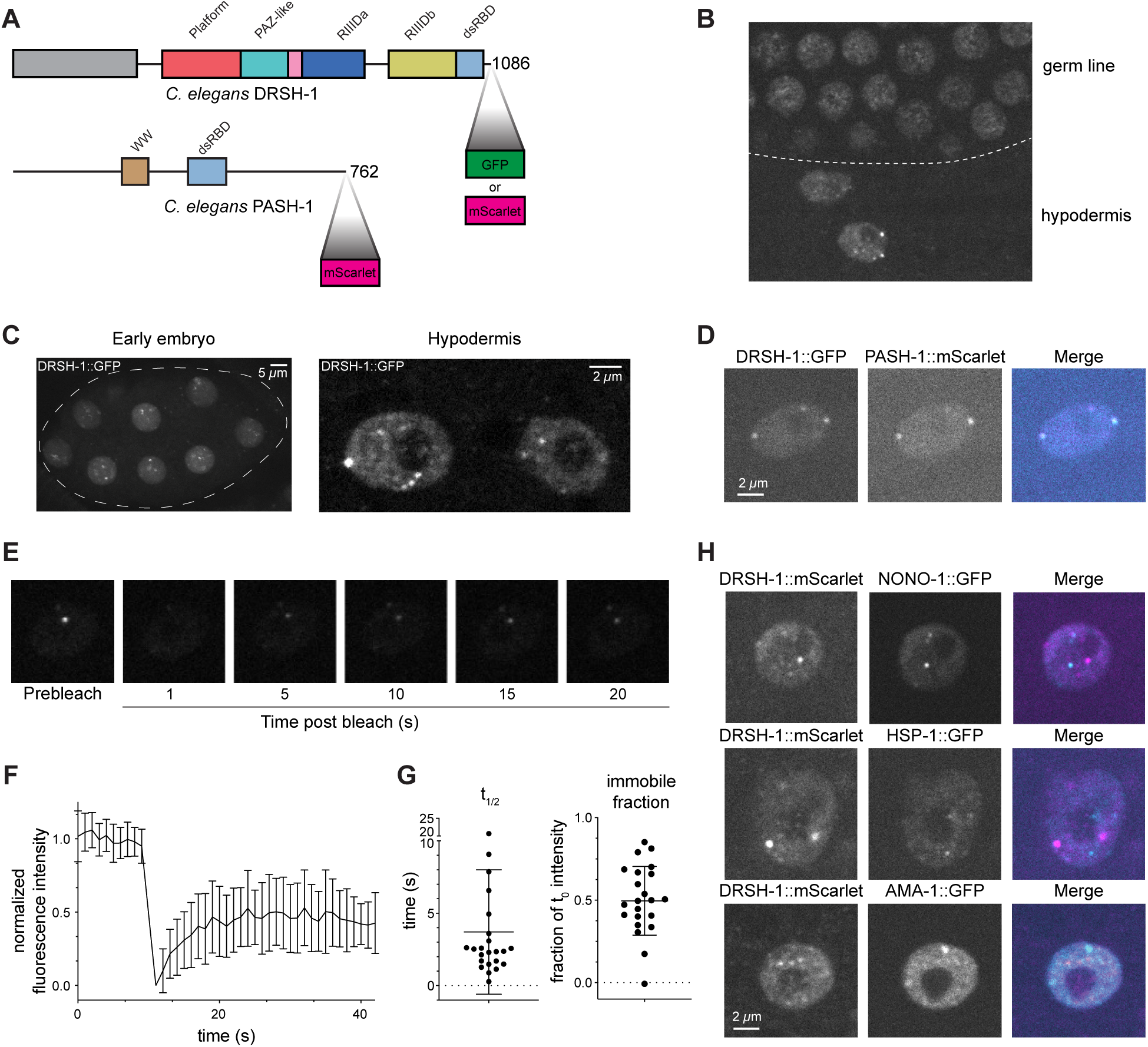
Microprocessor forms distinct nuclear granules in select tissues. (A) Domain architecture of *C. elegans* DRSH-1 and PASH-1. GFP and mScarlet were fused to DRSH-1 and PASH-1 at the C-terminus endogenously via CRISPR/Cas9. RIIID = RNase III domain, dsRBD = dsRNA binding domain. (B) Micrograph showing DRSH-1::GFP in germline nuclei lacking foci (top) and hypodermis nuclei containing foci (bottom). (C) Representative micrographs of DRSH-1::GFP in early embryos (left) and two hypodermal nuclei (right). Left: Dotted line outlines the embryo structure. Right: The large nucleoli characteristic of hypodermal nuclei exclude DRSH-1::GFP. Maximum intensity projection. (D) Representative images of co-localization of DRSH-1::GFP and PASH-1::mScarlet in a hypodermal cell. Single z plane shown. (E) Time-lapse images show fluorescence recovery of DRSH-1::GFP after a single focus is photobleached at 1.0 sec. Single z plane shown (F) FRAP recovery curves for DRSH-1::GFP. Data is reported as a mean +/- SD (n = 23 granules). (G) Half recovery time and immobile fraction were calculated based on fitted FRAP recovery curves for individual foci as shown in figure E. Bars are mean +/- SD. (H) Representative images show co-labeling of DRSH-1::GFP and known nuclear granule markers. Maximum intensity projections.

Next, we imaged live animals to characterize the distribution of Microprocessor. As expected, we observed DRSH-1::GFP localization primarily in the nucleus across different tissues (Figure 1B). In most tissues, the nuclear DRSH-1::GFP signal was diffuse (e.g. germ line in Figure 1B). Interestingly, we found prominent granule-like distribution of DRSH-1 in nuclei of early embryos and nuclei of both larval and adult hypodermis cells (Figure 1C). The diameter of these granules ranges from those at the diffraction limit (≤250nm) up to ∼1µm (Figure S2A). The proportion of DRSH-1::GFP that was localized to the granules was ∼2% of the total nuclear signal in the hypodermal cells (Figure S2B). To rule out that GFP contributes to granule formation, we also tagged DRSH-1 with highly monomeric mScarlet (Bindels et al., 2017) and consistently saw granule-like distribution in early embryo and larval and adult hypodermal nuclei (Figure S2C). Imaging of DRSH-1::mScarlet co-expressed with GFP::H2B shows the distribution of these granules throughout the nucleoplasm (Figure S2D). To determine whether the DRSH-1 granules contain both components of the Microprocessor, we generated a double tagged *drsh-1::gfp; pash-1::mscarlet* strain and observed strong co-localization of DRSH-1::GFP and PASH-1::mScarlet in these granules (Figure 1D). Taken together, the Microprocessor complex clusters in bright granule-like foci in specific tissues.

Given the granule-like distribution pattern, we wondered whether these granules are LLPS-mediated biomolecular condensates. To test this hypothesis, we performed fluorescence recovery after photobleaching (FRAP) experiments to bleach an entire focus and monitor fluorescence recovery by the influx of DRSH-1::GFP from the surrounding nucleoplasm. DRSH-1::GFP foci recovered quickly after bleaching (t_1/2_ = 4s ± 0.9s, SEM, n=23) with the intensity recovered to around 50% of the pre-bleaching level, indicating that the condensates consist of both a mobile fraction that exchanges with the nucleoplasmic fraction rapidly and also an immobile DRSH-1::GFP fraction that exchanges very slowly or not at all (Figure 1E-G). While this rapid FRAP is consistent with LLPS of DRSH-1::GFP foci, other mechanisms are also possible. Notably, while the use of live animals allows imaging of many tissues and discovery of tissue-specific foci, the *in vivo* system also limits our further characterization of the putative phase separation of the foci (such as via treatment with 1,6-hexanediol or biophysical approaches).

To determine if DRSH-1 granules coincide with known nuclear granules, we imaged fluorescently-tagged DRSH-1 alongside markers of nuclear granules: paraspeckles marked by NONO-1::GFP, nuclear stress granules labeled by HSF-1::GFP, RNA polymerase II transcriptional condensates marked by AMA-1::GFP, FUS granules labeled by FUST-1::GFP, splicing bodies marked by FOX-1::GFP, and heterochromatic foci marked by MET-2::mKate (Hills-Muckey et al., 2022; Pham et al., 2021). Although a previous report suggested that DGCR8 localizes to paraspeckles (Jiang et al., 2017), we failed to observe co-localization of either Microprocessor subunit with NONO-1 foci (Figure 1H and S2F). Furthermore, we did not observe any co-localization between DRSH-1 and other known nuclear granule markers (Figure 1H and S2E-F), with the exception of methyltransferase MET-2, where colocalization was only rarely observed: 17% (10 out of 59) of DRSH-1 granules colocalized with MET-2 (Figure S2G). In conclusion, we have found that the Microprocessor complex forms distinct nuclear foci in select tissues in *C. elegans* possibly through LLPS.

### DRSH-1 is a client protein in granule formation

Because we observed DRSH-1 granules that are distinct from known nuclear granules, we interrogated whether DRSH-1 itself is the driver of the putative LLPS or rather a client of a condensate formed by another scaffold molecule. If DRSH-1 is the driver of its own phase separation, then all DRSH-1 above the protein’s characteristic saturation concentration (c_sat_) would phase separate and the nucleoplasmic level would be at or below c_sat_. This would predict that tissues with observable DRSH-1::GFP foci have higher overall nuclear concentration of DRSH-1::GFP than tissues lacking DRSH-1::GFP foci. When we quantified the mean DRSH-1::GFP intensity in nuclei of hypodermis cells and adjacent germ cells (which lack DRSH-1::GFP foci), no significant difference of the average intensities was observed (Figure S2H-I). This observation suggests that granule formation is not solely due to DRSH-1::GFP protein concentration, and DRSH-1 is an LLPS client protein in hypodermal nuclei. However, because only ∼2% of DRSH-1::GFP resides in foci in hypodermal nuclei (Figure S2C), very small differences in concentration (as little as 2% higher in hypodermal nuclei) could underlie tissue specificity of granule formation; our detection method may not be sensitive enough to distinguish such small differences.

To further test whether granule formation depends on DRSH-1 concentration, we genetically controlled the concentration of DRSH-1. To this end, we engineered a situation in which the *drsh-1::gfp* allele is heterozygous with a *drsh-1* null allele (hemizygous), thus resulting in lower concentration of DRSH-1::GFP (Figure 2A). We then characterized the hemizygous strain in comparison to its homozygous *drsh-1::gfp* parental strain. In the hemizygous strain, we observed that the overall hypodermal nuclear signal of DRSH-1 was reduced to about 60% of the control (Figure 2B-C). If DRSH-1::GFP granule formation in the wild type setting is driven by DRSH-1::GFP exceeding its c_sat_ by ∼2%, then DRSH-1::GFP granules would be abolished in the hemizygous setting which reduces DRSH-1::GFP concentration well below the theoretical c_sat_. Instead, granules were still formed, though their intensity was reduced to a similar extent as the overall nuclear signal (Figure 2C-D). Reduced granule intensity in the hemizygous background held true when quantifying all foci together (Figure S3A) or when comparing the first or second brightest focus in each nucleus (Figure 2D, Figure S3A). (Intensity was the most accurate measurement of granule formation; we were unable to accurately measure granule size due the diffraction limit, and we were unable to precisely count granule number as many granules were not much brighter than the nucleoplasm, preventing accurate image thresholding.) Overall, DRSH-1::GFP does not obey a simple c_sat_ predictive model, and thus it likely is a client of other molecules in its recruitment to granules in specific tissues.

**Figure 2.**
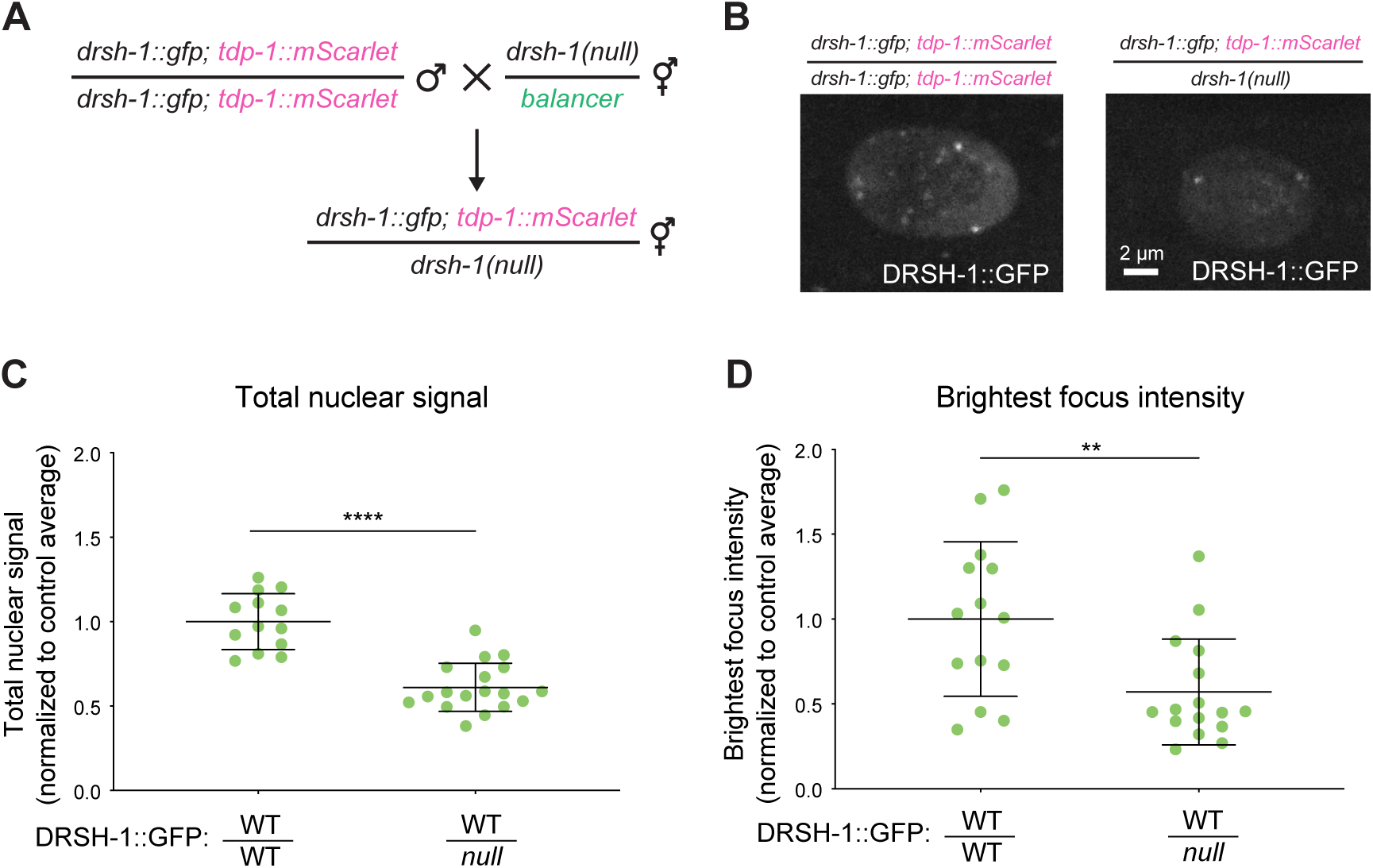
DRSH-1 granules are not strictly dependent on saturation concentration of DRSH-1. (A) Schematic of the genetic cross to generate *drsh-1::gfp/drsh-1(null)* heterozygous. *tdp-1::mScarlet* is a visual marker to differentiate cross progeny from self-progeny. Heterozygous worms were selected based on the visual marker. *drsh-1::gfp;tdp-1::mScarlet* homozygous worms were collected and used as control. (B) Representative images of DRSH-1::GFP in hypodermal cells in genetic background containing two (left) or one (right) allele of DRSH-1::GFP. Maximum intensity projection. (C) Quantification of the total nuclear signal of DRSH-1::GFP. (D) Quantification of the signal intensity of the brightest focus per nucleus across indicated genotypes in hypodermal cells. (C-D) N ≥ 6 animals per genotype Signal is normalized to the average signal in the control genotype (homozygous DRSH-1::GFP).

A previous study showed that expression of a large miRNA cluster attracted an immobile aggregate of Microprocessor to the genomic locus (Bellemer et al., 2012). To determine whether high expression of clustered miRNAs may also drive the tissue-specific Microprocessor condensate formation observed here, we examined the relative expression of clustered miRNAs in a condensate-forming (hypodermis) and non-condensate forming (neuron) tissue, based on tissue-specific miRNA sequencing (Wang et al., 2024). We observed no enrichment of clustered miRNA expression in hypodermis relative to neuron, providing no support for the possibility of clustered miRNA expression driving tissue-specific Microprocessor condensates (Figure S3C).

### Deletion of DRSH-1 IDRs reduces but does not abolish granule recruitment of DRSH-1

The N-terminal region of DROSHA contains intrinsically disordered regions and is proline-enriched (Son et al., 2023). Although the proline-enriched nature is not conserved in *C. elegans* DRSH-1, the disorderedness is conserved (Son et al., 2023). Numerous reports have established the significant contribution of IDRs to LLPS (Dignon et al., 2020); therefore we wondered whether the IDRs of DRSH-1 are important for mediating the recruitment of Microprocessor to granules. To understand the function of the N-terminus of DRSH-1 in its distribution and functionality, we utilized IUPred to predict disorderedness of the region and identified two putative disordered domains (Erdős & Dosztányi, 2020; Mészáros et al., 2018), namely IDR1 and IDR2 (Figure 3A). We used CRISPR to establish mutants in which IDR1, IDR2, or both IDRs are deleted, in both wild type and *drsh-1::gfp* backgrounds. Using the different IDR deletions in the *drsh-1::gfp* background, we assayed the expression of these truncated proteins through western blotting. Migration of the truncated proteins was consistent with their expected sizes. The mutations did not have significant effects on DRSH-1::GFP protein level in the whole animal samples (Figure 3B).

**Figure 3.**
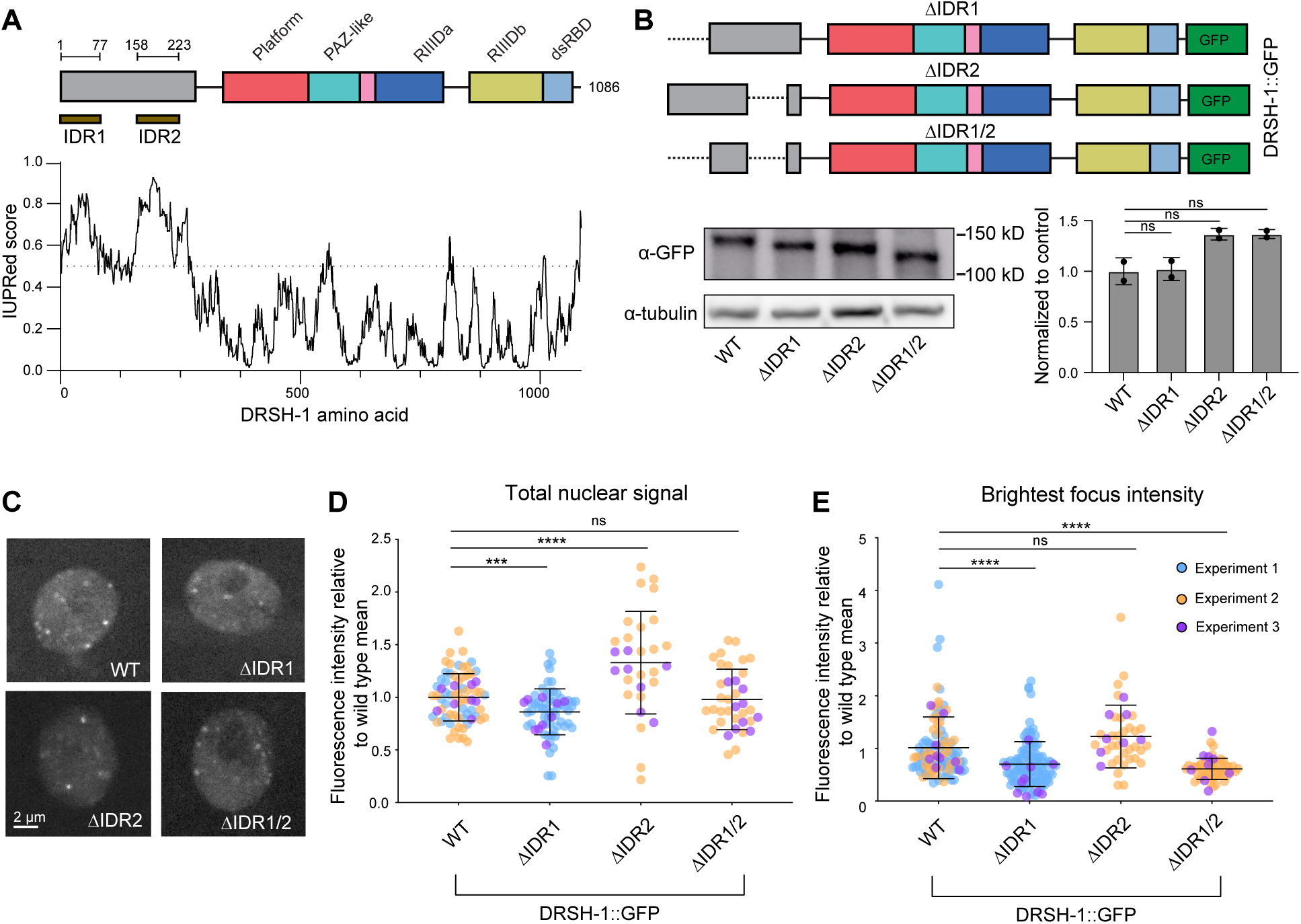
Deletion of N-terminal IDRs reduces but does not abolish DRSH-1 granule recruitment. (A) IUPRed was used to predict two intrinsically disordered domains: IDR1 (A.A. 1-77) and IDR2 (A.A. 158-233). (B) Schematics of IDR deletions generated in the *drsh-1::gfp* background using CRISPR/Cas9 and representative western blot images to show truncations of the DRSH-1::GFP fusion proteins in whole animal samples. Western quantification was normalized first to tubulin and then compared to wild type. Two tailed unpaired Student’s test: ns, not significant. (C) Representative images of DRSH-1::GFP signal in hypodermal nuclei across different genotypes. Maximum intensity projections. (D) Super plot showing the quantification of total nuclear signal of DRSH-1::GFP. (E) Super plot showing the quantification of the intensity of the brightest focus in each nucleus. (D-E) N ≥ 15 animals per genotype combining three experiments. Signal is normalized to the average signal in the control genotype (wild type DRSH-1::GFP) from the same imaging session. Data are color-coded by imaging session.

To understand the impact of the IDR deletions on the granule recruitment of DRSH-1::GFP, we imaged the hypodermal nuclei of each of the mutants and compared them to wild type. Overall nuclear signal was slightly reduced in ΔIDR1, slightly increased in ΔIDR2, and similar to wild type in the ΔIDR1/2 background (Figure 3D). (Although these results partially agree with trends observed by western blot, differences are likely due to the difference in sample type: whole animal samples with mixed tissues for western blot versus hypodermal tissue nuclei for imaging.) We found that ΔIDR1 results in slightly lower intensities of granules than wild type; this was the case either when quantifying all foci together (Figure S4A) or when comparing the first or second brightest focus in each nucleus (Figure 3E, S4B). If the IDR1 has a direct role in DRSH-1 recruitment to granules, then its effect on granule intensity should be greater than that of the hemizygous genotype, which had a similar impact as ΔIDR1 on overall nuclear signal (reducing it to 60% of wild type). However, the reduction in granule intensity was comparable to that in the hemizygous background (Figure 3E, Figure 2D), suggesting that the IDR1 itself has no impact on granule recruitment beyond its effect on DRSH-1::GFP concentration. The ΔIDR2 mutation alone did not significantly affect the intensity of all foci quantified together (Figure S4A) and had inconsistent impacts on the intensity of the two brightest granules per nucleus (Figure 3E, Figure S4B). Finally, the ΔIDR1/2 combined mutation reduced granule intensity without affecting overall nuclear DRSH-1::GFP signal (Figure 3D-E, Figure S4). Because the combined mutation – but not the single IDR deletions – decouples granule intensity from DRSH-1::GFP level (reducing intensity without affecting level), this suggests that the two IDRs may act redundantly to promote granule recruitment. This reduction in granule intensity is again mild, indicating that the IDRs are not strictly required for granule recruitment of DRSH-1::GFP, and other parts of the protein also likely contribute. Overall, the IDR mutations had moderate impacts on granule formation, which in the case of ΔIDR1 reflected its impacts on DRSH-1::GFP level.

### An IDR is important for biogenesis of a subset of miRNAs in *C. elegans*

Despite the modest impact of individual IDR mutations on DRSH-1 level and granule recruitment, we observed a strong impact on development in ΔIDR1 mutant animals. The ΔIDR1 but not ΔIDR2 animals showed bursting (vulval rupture) with 9% penetrance, while ΔIDR2 animals were superficially wild type (Figure 4A and 4B). Similar results were obtained when the deletions were introduced in wild type or *drsh-1::gfp* backgrounds, confirming not only the phenotype but also the functionality of the *gfp* tagged DRSH-1 protein (Figure 4B). No other phenotypes were observed; non-bursting ΔIDR1 animals exhibited wild type egg retention, brood size and embryonic viability (Figure S5A-C). The ΔIDR1/2 double mutant mirrored the ΔIDR1 phenotype with 9% penetrance of bursting, indicating that IDR1 and IDR2 do not act redundantly with respect to this aspect of development (Figure 4B).

**Figure 4.**
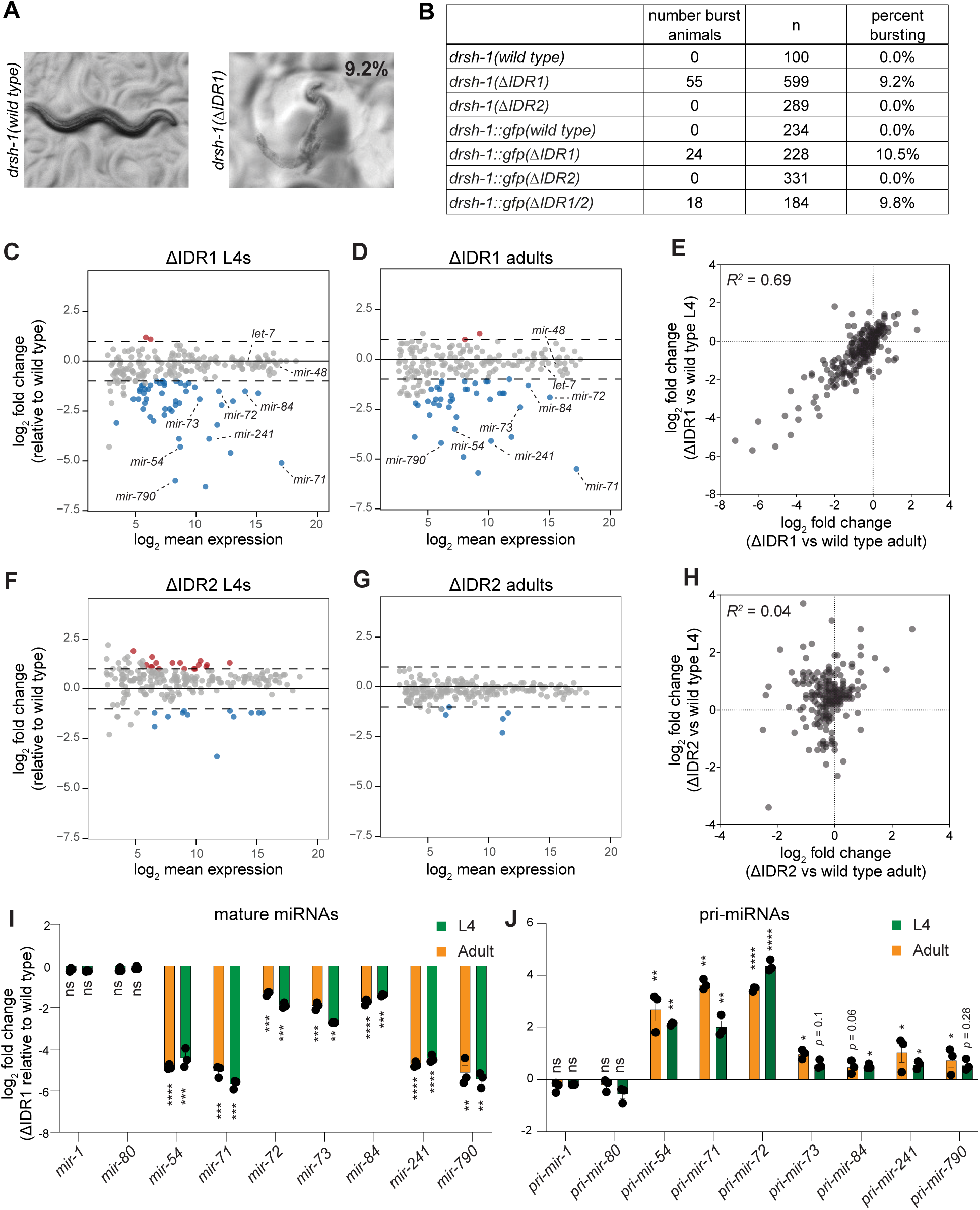
DRSH-1 IDR1 is important for the biogenesis of a subset of miRNAs. (A) Representative images showing the bursting phenotypes observed in 9.2% of ΔIDR1 young adult animals. (B) Table listing the penetrance of the bursting phenotype in different genotypes. IDR deletions were generated in both wild type and *drsh-1::gfp* backgrounds. (C-D, F-G) MA plots showing average abundance on the X-axis and the log_2_fold change on the Y-axis. Sequencing analysis results were obtained from DESeq2 using total piRNA reads for normalization. miRNAs showing significant changes compared to wild type of the same stage are shown in blue (downregulated, log_2_fold change < −1 and *p*_adj_ < 0.05) and red (upregulated, log_2_fold change > 1 and *p*_adj_ < 0.05. (C and D) MA plots showing miRNA abundance changes in ΔIDR1 versus wild type for L4 and adult, respectively (E) X-Y plot showing the correlation of log_2_fold change for ΔIDR1/wild type between L4 (Y-axis) and adult (X-axis). (F and G) MA plots showing miRNA abundance changes in ΔIDR2 versus wild type for L4 and adult, respectively. (H) X-Y plot showing the correlation of log_2_fold change for ΔIDR2/wild type between L4 and adult samples. (I) Relative levels of mature miRNAs as measured by Taqman qPCR in ΔIDR1 L4 and adults, normalized to wild type of the same stage. (J) Relative levels of primary miRNA transcripts as measured by qRT-PCR in ΔIDR1 L4 and adults, normalized to wild type of the same stage. (I-J) *Sn2429* and *gpd-1* were used as internal controls for mature miRNAs and pri-miRNAs, respectively. Two tailed unpaired Student’s test: (*) P < 0.05, (**) P < 0.01, (***) P < 0.001, (****) P < 0.0001.

The vulval bursting phenotype we observed is reminiscent of loss-of-function mutations in the miRNA pathway (Grishok et al., 2001; Reinhart et al., 2000). To examine the impact of the IDRs on miRNA biogenesis, we performed small RNA sequencing of the ΔIDR1 and ΔIDR2 mutants at the fourth larval (L4) and adult stages. Sequencing analysis using either total piRNAs or spike-ins as a normalization method showed very similar differential expression results (Figure 4 and S5D, Table S5-S12). Among a total of ∼200 miRNAs detected by sequencing (DEseq2 normalized mean reads > 5), 49 miRNAs were significantly downregulated in ΔIDR1 in comparison to wild type (DEseq2 log_2_FC < −1, *p*_adj_ < 0.05) at the L4 stage, and only two miRNAs were upregulated (log_2_FC > 1, *p*_adj_ < 0.05) (Figure 4C, Table S5). Moreover, we observed a very similar pattern of expression changes in ΔIDR1 animals compared to wild type at the adult stage, with 41 miRNAs downregulated and two upregulated (Figure 4D, Table S6). Changes in miRNA levels caused by ΔIDR1 were highly correlated between the two stages (Figure 4E), with 80% of the downregulated miRNAs in common between L4 and adult (Fisher’s exact test *p* < 1×10^-17^). Thus the ΔIDR1 mutation causes a consistent selective defect in miRNA production affecting the same set of miRNAs across life stages. Among the miRNAs downregulated at both stages are two members of the *let-7* family*, mir-84* and *mir-241*; the downregulation of these miRNAs may be the molecular basis of the bursting phenotype, which is reminiscent of *let-7* family loss of function mutants.

Unlike ΔIDR1 mutants, only minor changes in miRNA profile were observed in the ΔIDR2 mutants, with modest amplitude of changes and poor consistency between life stages (Figure 4F-H, Table S7-S8). In conclusion, IDR1 deletion results in a strong, selective defect in miRNA production across different life stages, whereas IDR2 deletion has little impact.

Next, we sought to determine whether the miRNA abundance changes we observed in ΔIDR1 animals are due to Microprocessor defects rather than other indirect effects. We first validated the downregulation that was observed in deep sequencing for seven mature miRNAs by qPCR (Figure 4I). We also included two miRNAs (*mir-1*, *mir-80*) as controls that were not differentially expressed in the sequencing data, nor were they altered in ΔIDR1 mutants by qPCR (Figure 4I). If the prominent downregulation of select miRNAs observed in ΔIDR1 animals is due to a defect in Microprocessor activity, then decreased mature miRNA levels should be accompanied by increased pri-miRNA levels. We measured pri-miRNA levels by qPCR and observed that this is the case: in all examined cases in which the mature miRNA decreased in ΔIDR1, the corresponding pri-miRNA level increased (Figure 4J). Furthermore, the examined mature miRNAs that were not sensitive to ΔIDR1 (*mir-1, mir-80*) also showed no changes in pri-miRNA level (Figure 4J). Overall, this suggests that DRSH-1 IDR1 is crucial for the efficient Microprocessing of a select subset of miRNAs.

Importantly, neither the bursting phenotype nor the molecular phenotype were due to IDR1’s effect on DRSH-1 level or granule recruitment, since neither phenotype was observed in the DRSH-1::GFP hemizygous setting which displayed similar reduction in DRSH-1::GFP levels and granule recruitment as in ΔIDR1 (Figure 2). Small RNA sequencing revealed that the 60% dosage of DRSH-1::GFP and resulting lower granule intensity in the DRSH-1::GFP hemizygous setting did not result in significant changes in miRNA expression profile (Figure S3B, Table S15). Moreover, the hemizygous worms were superficially wild type without any noticeable phenotypes (no bursting). Therefore, the effects of the ΔIDR1 mutation are not recapitulated by orthogonal means of reducing protein dosage and granule recruitment, demonstrating instead that the IDR1 plays a specific role in Microprocessor-mediated cleavage of select substrates.

### IDRs have minimal impacts on cleavage site fidelity, except for *mir-259*

We next examined the impact of IDR deletions on cleavage precision by Microprocessor. While the small RNA sequencing method used above (NEBNext) allows for accurate comparison of miRNA expression levels between genotypes, this method suffers from sequence-dependent bias in adaptor ligation efficiency, which confounds comparison of different miRNAs or miRNA isomers (isomiRs) to each other. Therefore, to assess cleavage precision, we performed bias-minimized small RNA sequencing, in which degenerate bases at the ligated end of the library construction adaptors allow for equalized cloning efficiency of different miRNA sequences (H. Kim et al., 2019; Mandlbauer et al., 2024). As expected, the bias-minimized method showed very similar differential expression results when comparing genotypes (ΔIDR1 or ΔIDR2 adults to wild type) (Figure S5E, Table S13-S14).

Next, we globally examined the cleavage fidelity in wild type and ΔIDR1 or ΔIDR2. We divided the analysis into 5p-derived miRNA strands – of which Microprocessor-mediated cleavage generates the 5’ end – and 3p strands, of which Microprocessor cleaves the 3’ end (Kwon et al., 2019a) (Figure 5A). Apparent fidelity of Microprocessor cleavage (% with the canonical end position, set as position 0) was higher for 5p strands than 3p strands (Figure 5B, Figure S6). However, miRNA 3’ ends (the position assessed for the 3p strands) are known to undergo remodeling that includes exonucleolytic trimming and enzymatic tailing, which cannot be distinguished from Microprocessor cleavage imprecision if the trimmed/tailed isomer still fully matches the genome-encoded sequence. Therefore, we interpret the greater degeneracy in 3p strand 3’ end position to reflect tailed and trimmed isoforms. Next, we focused on the miRNAs that are downregulated in ΔIDR1. These miRNAs did not display significant differences in the position of their Microprocessor-cleaved ends, with the exception of *mir-54*, suggesting that their downregulation is due to loss of Microprocessor cleavage efficiency, not loss of precision (Figure 5C). Since *mir-54* is a 3p-derived miRNA, the shift in its 3’ end position could be due to a shift in Microprocessor cleavage position or due to more prevalent terminal uridylation, the most common form of miRNA tailing in *C. elegans* (Vieux et al., 2021).

**Figure 5.**
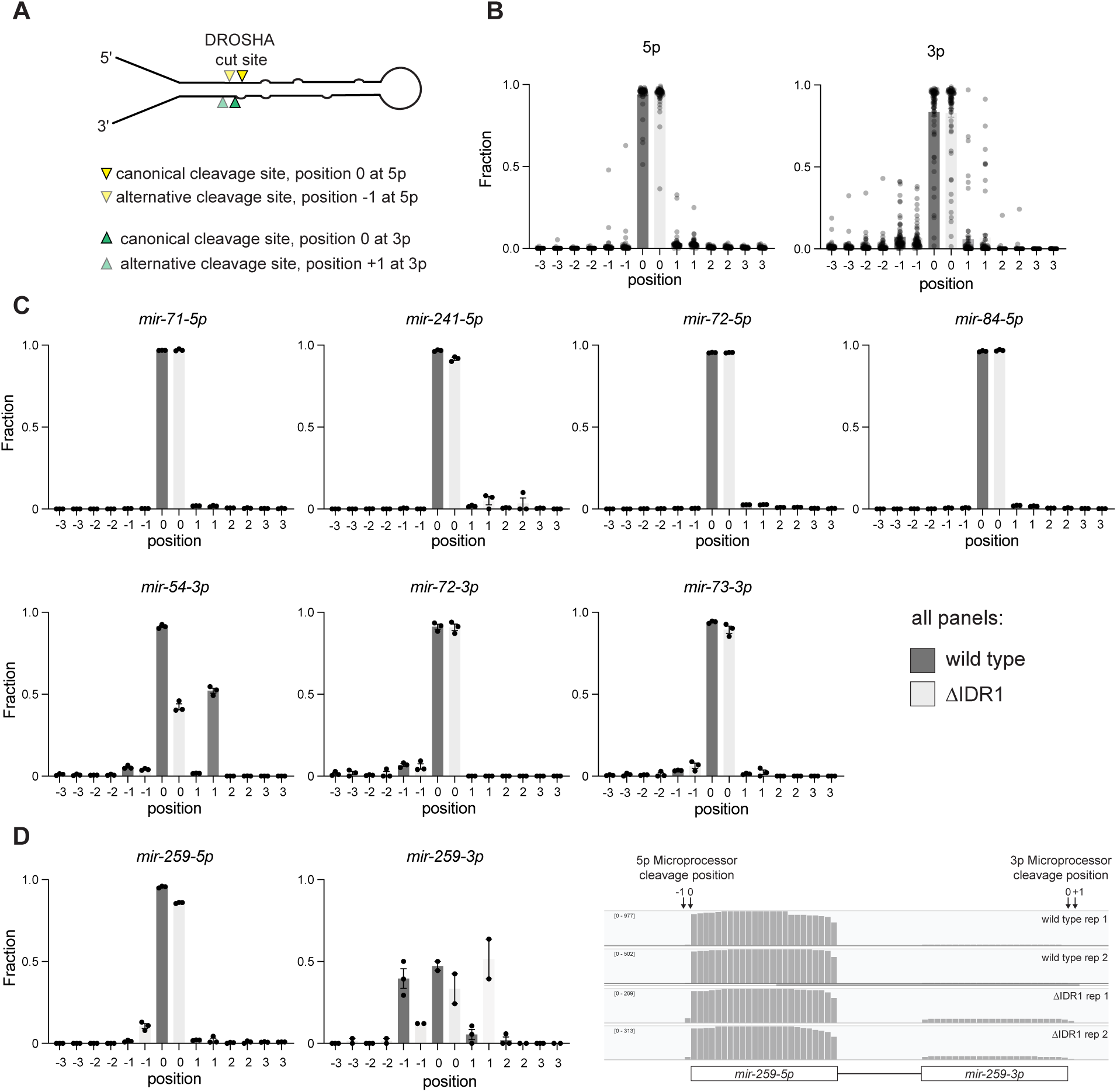
DRSH-1 IDR1 has almost no effect on Microprocessor precision. (A) Illustration of pri-miRNA showing an example of relative positions of canonical and alternative Microprocessor cleavage sites. (B) Distribution of terminal miRNA positions for 5’ ends of 5p miRNA strands and s of 3p strands in wild type and *drsh-(1*Δ*IDR1)* animals at adult stage. (C) Examples of 5p and 3p strand isomiR distribution of individual miRNAs. These miRNAs are downregulated significantly in abundance in the *drsh-(1*Δ*IDR1)* animals. (D) isomiR distribution of both strands of *mir-259* showing decreased precision in the Δ*IDR1* background.

We next extended our analysis to all miRNAs by comparing the cleavage fidelity (% canonical cleavage site) between wild type and ΔIDR1 or ΔIDR2. Only one miRNA showed a significant decrease in cleavage fidelity, *mir-259*, which showed shifted end positions in ΔIDR1. Both strands of *mir-259* were affected, suggesting a greater frequency of Microprocessor cleavage shifted 1bp toward the basal junction relative to the canonical cleavage site in ΔIDR1 (Figure 5A, C). While future work may elucidate what renders the pri-*mir-259* cleavage fidelity sensitive to IDR1, it remains the exception, and generally the impact of IDR1 on miRNAs is solely through changes in abundance, not cleavage position.

### *Cis* sequences of primary *mir-241* confer sensitivity to IDR1 deletion

We observed that ΔIDR1 mutation in DRSH-1 induces a consistent defect in Microprocessing affecting a subset of pri-miRNA substrates; we therefore sought to understand what underlies the specificity of dependence upon the IDR1 for biogenesis. To this end, we tested the association between various features of pri-miRNAs and their sensitivity to the IDR1 deletion (Figure 6A).

**Figure 6.**
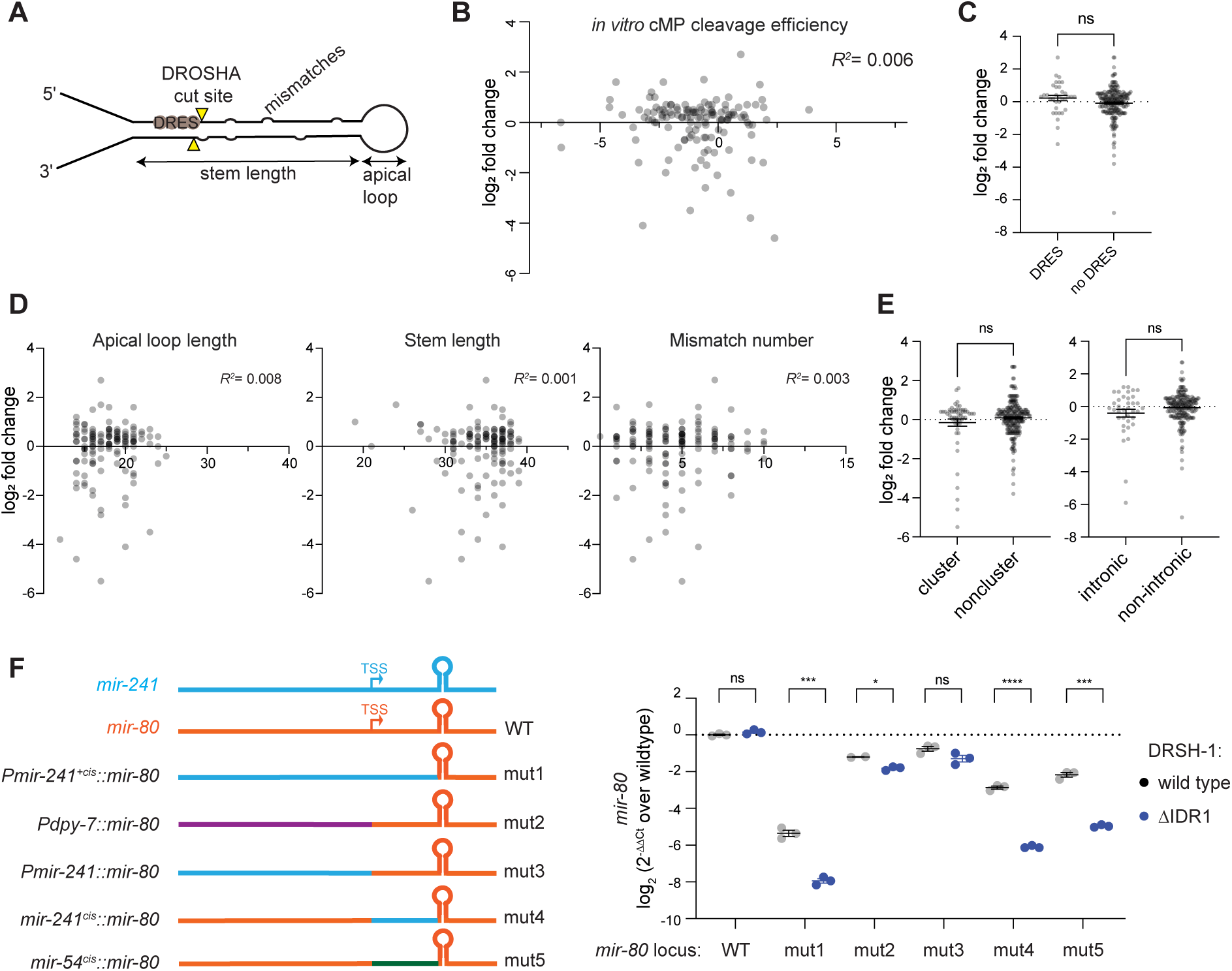
5’ flanking sequence of miRNA confers IDR1-sensitivity. (A) Schematic diagram of local features of a pri-miRNA hairpin. (B) X-Y plot showing correlation between IDR1 sensitivity measured by miRNA log_2_ fold change (ΔIDR1/wild type L4) on the Y-axis and *in vitro C. elegans* Microprocessor cleavage efficiency from Nguyen, et al. 2023 on the X-axis. (C, E) IDR1 sensitivity between miRNA groups based on classifications: containing DROSHA dsRNA recognition sites (DRES) site or not in (C), clustering or standalone and intronic or exonic in (E). Two tailed unpaired Student’s t-test, ns, not significant. (D) X-Y plots showing correlation between IDR1 sensitivity, and different predicted pri-miRNA secondary features, including apical loop length, stem length and mismatch number. (F) Left: Schematics showing genome edits made at the *mir-80* locus. Genomic structure of *pri-mir-241* is shown in blue, and orange shows the genomic structure of *pri-mir-80*. Hairpin structure depicts the pri-miRNA hairpin defined by the basal junctions. TSS denotes transcription start site. Right: *mir-80* Taqman qPCR comparing different mutants in DRSH-1 wild type or Δ IDR1 background at adult stage. Two-tailed unpaired Student’s t-test: (ns) not significant, (*) P < 0.05, (**) P < 0.01, (***) P <0.001, (****) P < 0.0001.

Although multiple sequence motifs associated with optimal Microprocessor primary miRNA substrates are well annotated in mammalian cells (Auyeung et al., 2013; Fang & Bartel, 2015; Jin et al., 2020; Kwon et al., 2016, 2019b; T. A. Nguyen et al., 2015; Partin et al., 2017), most of these motifs are not enriched in *C. elegans* pri-miRNAs, with the exception of the midBMW and mGHG motifs (Kwon et al., 2019b; Li et al., 2020, 2021), neither of which predicted dependency on IDR1 (Figure S7A-B). A recent effort to characterize the *C. elegans* Microprocessor *in vitro* has advanced our understanding of its cleavage parameters and identified a DROSHA <u>d</u>sRNA <u>re</u>cognition <u>s</u>ites (DRES) motif that correlates with substrate optimality (T. L. Nguyen, Nguyen, Ngo, & Nguyen, 2023; T. L. Nguyen, Nguyen, Ngo, Le, et al., 2023). We examined whether the *in vitro* Microprocessor cleavage score or the DRES score were correlated with miRNA sensitivity to ΔIDR1 and again found no association (Figure 6B-C, S7C).

We then analyzed the features of primary miRNA secondary structure predicted by miRGeneDB (Clarke et al., 2025), including apical loop length, stem length, mismatch count, lower stem length, and bulges. However, no significant correlation was observed between any of these structural features and ΔIDR1 sensitivity (Figure 6D and S7D-E). Additionally, we considered the genomic contexts of pri-miRNAs, including clustering status and intronic/exonic position that were previously reported to be important for the processing of certain subsets of miRNAs (Donayo et al., 2019; Fang & Bartel, 2020; Hutter et al., 2020; Janas et al., 2011; Lataniotis et al., 2017; Pawlicki & Steitz, 2008; Shang et al., 2020; Slezak-Prochazka et al., 2013; Thivierge et al., 2024; Truscott et al., 2016). Moreover, the N-terminus proline-enriched domain of human DROSHA has been shown to mediate intronic miRNA biogenesis (Son et al., 2023). However, our analysis did not show any significant difference in ΔIDR1 sensitivity between these miRNA categories (Figure 6E).

At the organismal level, miRNA expression is known to be spatially regulated and often exhibit tissue and cell-type specificity (Aboobaker et al., 2005; Alberti et al., 2018; Brosnan et al., 2021; Landgraf et al., 2007; Martinez et al., 2008). Given that we observed tissue-specific formation of granules and obvious developmental phenotypes related to hypodermal cells, we wondered whether tissue of expression could contribute to ΔIDR1 sensitivity. To test this hypothesis, we replaced the upstream sequence of an ΔIDR1-insensitive miRNA (*mir-80*) with the upstream sequences of an ΔIDR1-sensitive miRNA (*mir-241*). Intriguingly, we noticed that this ∼2kb fragment conferred ΔIDR1 sensitivity to *mir-80* (Figure 6F, mut1). To determine whether this fragment conferred sensitivity by driving *mir-80* expression in the hypodermis, we then replaced the sequence upstream of *mir-80* with a known hypodermis promoter (*dpy-7*), but this did not confer sensitivity to ΔIDR1 to *mir-80* to the same extent as the 2kb *mir-241* upstream fragment (Figure 6F, mut2). We thus revisited the 2kb sequence and divided it into the 1.7kb fragment upstream of the pri-*mir-241* TSS (R. A.-J. Chen et al., 2013; Gu et al., 2012; Kruesi et al., 2013; Saito et al., 2013) and the remaining 293bp that lies between the TSS and the predicted basal junction of pri*-mir-241*. We then replaced the corresponding fragments in the *mir-80* locus with the fragments of the *mir-241* locus. Strikingly, the 293bp cis element, but not the 1.7kb upstream fragment conferred sensitivity of *mir-80* to ΔIDR1 (Figure 5F, mut4 and mut3), similar to the full 2kb fragment of *mir-241*. To confirm that the 5’ flanking sequence is conferring IDR1 sensitivity, we replaced the 293bp fragment of *mir-80* with one from another IDR-sensitive miRNA, *mir-54,* and again observed conferral of IDR1-dependence (Figure 6F, mut5). To determine whether the sequence swaps interfered with the canonical cleavage of *mir-80* in the variant genetic backgrounds, we prepared small RNA libraries using the bias-minimized cloning method. This analysis recapitulated the *mir-80* abundance changes detected through miRNA Taqman qPCR (Figure S8A), and the cleavage patterns were highly canonical for all *mir-80* variant loci, suggesting that – similar to wild type IDR1-sensitive loci – the IDR1-sensitive variants of pri-*mir-80* rely on IDR1 for efficiency, not precision, of Microprocessor-mediated cleavage (Figure S8B). Together, this suggests that flanking sequences near the base of the pri-miRNA hairpin determine dependence on DRSH-1 IDR1 for maximally efficient processing.

## Discussion

In this study, we describe the formation of distinct nuclear DRSH-1 foci in select tissues in *C. elegans*, consistent with recruitment to LLPS condensates. Although a previous report hinted at the potential co-localization between DGCR8 and paraspeckles (Jiang et al., 2017), our study provides the first evidence of Microprocessor condensates in animals. We demonstrate that DRSH-1 is a client protein in condensate formation since the simple phase separation model defined by >*c_sat_* of DRSH-1 is incompatible with the behavior of these granules. What scaffolds Microprocessor foci and governs their tissue specificity will be a crucial question as more components of these granules are identified. The current study is also limited by the diffraction limit (∼250nm) of our imaging methods; further characterization of the granules by super resolution microscopy will provide finer details of the condensates’ structure and potentially identify smaller-scale clusters in tissues other than embryos and hypodermal cells. Which regions of DRSH-1 itself are required for its recruitment to granules is another open question; the N-terminal IDRs redundantly promote DRSH-1 granule recruitment, but their impact is modest, suggesting other regions of the protein also support recruitment.

The ability of Microprocessor to be recruited to condensates may provide a new layer of Microprocessor regulation that was not previously explored. A lack of conditions in which granule formation is completely abolished limits our ability to assess their function. One possibility is that the Microprocessor condensates are “incidental condensates” resulting from the biophysical properties of the complex and the nuclear context but not providing functionality beyond that of diffuse Microprocessor (Putnam et al., 2023). Understanding the factors involved in scaffolding of Microprocessor-containing granules will aid in perturbing these foci to probe their function.

Whereas the DRSH-1 IDRs had a modest role in Microprocessor recruitment to granules and modest effects on DRSH-1 concentration, IDR1 clearly contributes to Microprocessor function. The ΔIDR1 mutants exhibited strong molecular and developmental phenotypes related to Microprocessing defects.

The hypodermis is both a site of apparent Microprocessor condensates and the tissue of origin of the ΔIDR1 mutant phenotype. This led us to interrogate the relationship of these phenomena. Using genetics, especially the implications of the DRSH-1 hemizygous setting, we were able to separate the IDR1 function in biogenesis from DRSH-1 focus formation. The DRSH-1 hemizygous setting demonstrates that enforcing a modest reduction in DRSH-1 concentration and focus formation – comparable to those observed in the ΔIDR1 background – does not necessarily lead to defects in Microprocessing. We propose that careful modulation of protein concentration can be a general means for avoiding guilt-by-association between specific IDR-driven molecular processes and co-occurring LLPS.

We have shown that DRSH-1 IDR1 strongly contributes to biogenesis of a subset of miRNAs, independent of known features of *C. elegans* Microprocessor substrate optimality. The 5ꞌ flanking region of two IDR1-sensitive miRNAs each conferred sensitivity to an IDR1-insensitive miRNA; these interesting observations suggest that a cis-acting element in this region of the pri-miRNA could be interacting with the IDR1 domain. The fact that miRNAs derived from the same primary transcript behaved differentially in response to IDR1 deletion – i.e. *mir-54* is significantly IDR1-sensitive, whereas both *mir-55* and *mir-56* did not respond to IDR1 deletion (Figure S9) – further supports the model that a cis element very close to the miRNA hairpin could mediate IDR1-dependence. So far, our attempts at motif finding and structural prediction have not yielded candidate cis elements whose functionality can be tested. Our findings are interestingly similar to the involvement of cis elements in determining pri-miRNA reliance on the N-terminal IDRs of human DROSHA, suggesting that these sequence-divergent regions of orthologous proteins may maintain conserved function (Son et al., 2023). Future studies using high-throughput functional genomics should reveal what properties define the sequence or structural feature that is recognized by the IDR domain and whether the recognition is direct or indirect through other RNA binding proteins (Figure 7).

**Figure 7.**
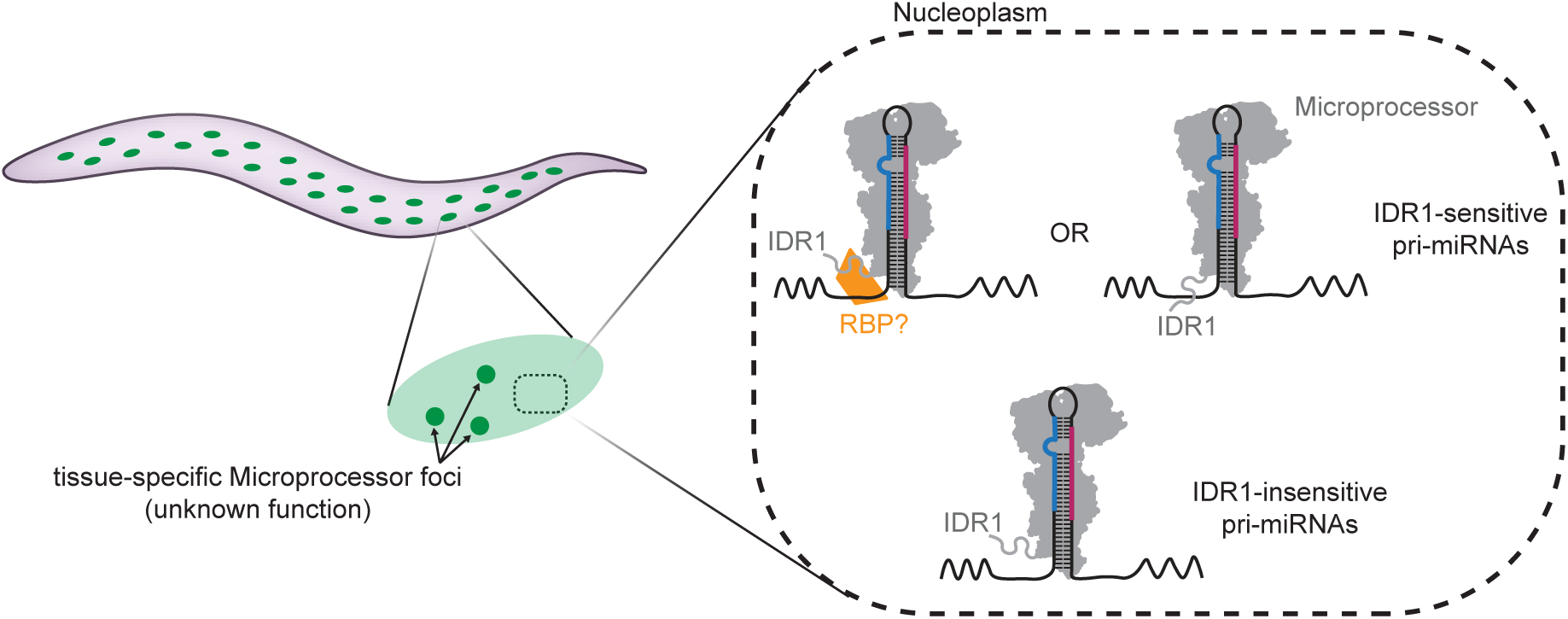
Model of DRSH-1 IDR1 function. Left: Microprocessor forms distinct foci in select tissues, such as hypodermis. Right: DRSH-1 IDR1 functions in miRNA maturation independent of Microprocessor focus formation. Sequences flanking the pri-miRNA hairpin confer sensitivity to the presence of IDR1. IDR1 may interact directly or indirectly with sequences flanking the pri-miRNA hairpin to promote Microprocessing.

## Materials and Methods

### C. elegans culture

*C. elegans* were cultured using standard protocols at 20°C (Stiernagle, 2006). L4 and adult worm samples were collected 48 h and 60 h after L1 synchronization, respectively.

### Genome editing

Unless otherwise mentioned, strains generated in this study were engineered endogenously using CRISPR-Cas9. CRISPR-Cas9 mediated genome editing is performed as previously reported (Yang et al., 2020). In brief, IDT alt-R guide RNAs were annealed by mixing crRNA and tracrRNA to a final concentration of 10µM each using IDT duplex buffer, which was then incubated at 95°C for 5 min followed by gradual temperature decline to room temperature. Repair templates provided for CRISPR were column purified after PCR synthesis. Final injection mix contains annealed gRNA at final concentration of 1µM, IDT Cas9 at final concentration of 2µM and repair template at final concentration ∼200-400ng/µl (Paix et al., 2014). A gRNA targeting *dpy-10* was used as a co-CRISPR marker for selection (Arribere et al., 2014). After injection, Dpy animals were cloned and singled as F1s. F2 animals were then genotyped for desired edits.

GFP donor fragments were amplified using plasmid pCM1.35 as a template and primers containing ∼35-55 bp homology arms (Merritt et al., 2008; Paix et al., 2014). mScarlet donor molecules were amplified using plasmid pGLOW30 as a template and primers containing ∼35-55 bp homology arms (Witten et al., 2023). Promoter sequences of *mir-241*, *mir-84* and *dpy-7* were amplified from wild type genomic DNA using primers listed in table S2. Transcription start sites were determined using the Wormbase TSS database (R. A.-J. Chen et al., 2013; Gu et al., 2012; Kruesi et al., 2013; Saito et al., 2013).

### Western Blotting

Three 10cm NGM plates seeded with 2,000 L1s were cultured at 20°C for 60 h, until animals reached adulthood. Animals were then rinsed off the NGM plates with M9 buffer and washed three times with M9 before being pelleted and snap frozen. Upon thawing, the worm pellets were mixed with equal volume of cold 2x worm lysis buffer (60 mM HEPES pH 7.4, 100 mM KCl, 0.4% Triton X-100, 4 mM MgCl_2_, 20% glycerol, 4 mM DTT, 2× cOmplete, Mini, EDTA-free Protease Inhibitor Cocktail) and lysed using a Bioruptor Pico (Diagenode) for 10 cycles (30 seconds on and 30 seconds off). Crude worm lysate was then cleared by centrifugation at 16,000x*g* for 10 min. Protein concentration of the resulting supernatant was then quantified using BCA assay. Equal amounts of protein were loaded onto a 4-15% gradient Tris Glycine gel (Biorad). Membranes were then probed with anti-GFP antibody (Roche) at 1:1,000 and anti-tubulin antibody DM1A (Abcam) at 1:1,000.

### Immunoprecipitation

Worms were harvested from 10cm NGM plates and washed three times with M9 buffer. Worm pellet was flash frozen on dry ice and stored in −80°C until use. Upon thawing, equal volume of 2X IP lysis buffer (100mM Tris-HCl, 200 mM KCl, 5mM MgCl2, 0.2% NP-40, 1mM PMSF, 20% glycerol) supplemented with proteinase inhibitor cocktail was added to the worm pellet. Worm lysis was carried out using Bioruptor sonicator with 7 cycles of 30 seconds ON and 30 seconds OFF. Worm lysate was then cleared by centrifugation at 15,000 rpm for 10 minutes. The supernatant was transferred to a new tube and used as input for immunoprecipitation. Proteintech GFP-trap agarose beads (10µl) were added to 2mg of input lysate and incubated at 4°C for 2 hours on a rotator. Beads were then collected by centrifugation at 1,000rpm for 1 minute, and washed three times with 1X IP lysis buffer. 2X SDS Laemmli sample buffer was added to the beads and incubated at 95°C for 5 minutes. Western blot was performed as above using anti-GFP antibody (Proteintech anti-GFP PABG1) and anti-RFP antibody (Proteintech anti-RFP 6G6).

### IsomiR analysis

To analyze isomiR distribution, aligned bam files were converted to fastq and collapsed to tabular files containing read sequence and count information. SeqBuster was then used to annotate miRNAs/isomiRs with the following parameters -sub 0 -trim 3 -add 3 -s cel. Annotation information from miRbase was used as a reference file for seqBuster (Pantano et al., 2010). A custom script was developed to analyze the distribution of isomiRs potentially resulting from variation of Microprocessor cleavage sites. Briefly, miRNAs with baseMean > 25 from DEseq2 analysis were included in this analysis. To minimize the confounding effects of miRNA tailing on isomiR distribution, miRNA reads with unambiguous untemplated additions at the 3’ end were removed from the analysis.

### Microscopy

Fluorescent microscopy was performed using a Nikon Eclipse Ti2 microscope (Nikon Instruments, Melville, NY, USA) equipped with a W1 spinning disk confocal head (Yokogawa, Life Science, Tokyo, Japan), 405-, 488-, 561-, and 641-nm laser lines, a 100x/1.49 total internal reflection fluorescence oil immersion objective (Nikon Instruments) and a Prime BSI cMOS camera (Teledyne Photometrics, Tuscon, AZ, USA). Fluorescence recovery after photobleaching (FRAP) was performed using an Opti-MicroScan unit with a 405 nm laser. Nikon Elements software was used to control the microscope and to collect images (Nikon Instruments). Image analysis was performed using FIJI (Image J; National Institutes of Health, Bethesda, MD, USA).

### FRAP

FRAP experiments were performed using the microscope described above. Hypodermal cells nearest the coverslip were selected for imaging in which a single confocal plane was acquired using 2×2 binning. One prominent fluorescent focus was selected for FRAP using a 10 millisecond, 405 nm laser exposure, which was sufficient to partially photobleach the focus. Movies of a single focal plane using the 488nm laser were then collected using 2×2 binning and one image/second. Only movies where the partially photobleached focus stayed within the imaging plane were used for analysis.

For analysis, total integrated fluorescence measurements of a region of interest including the bleached focus were recorded at each timepoint. Background subtraction was performed using a nuclear region of identical area adjacent to the focus. Correction for photobleaching resulting from timelapse image acquisition using the 488nm laser was performed by using the decrease in fluorescence across the entire image. Data was normalized by setting the average focus intensity from 10 pre-bleach frames to 1, and the first post-bleach frame to 0. The frame containing the bleaching laser pulse was excluded from the analysis. Figure 1F presents the average of all recovery curves ± standard deviation with time 0 equal to 10 frames prior to bleaching. The t_1/2_ of recovery and immobile fraction from individual recovery curves were determined following fitting (one phase association) using Prism (GraphPad, Boston, MA, USA).

### Fluorescence intensity measurements in hypodermal nuclei

Confocal Z-stack images of hypodermal cells close to the coverslip in live worms were acquired using the microscope described above. Only nuclei where the animal and the foci did not move substantially during acquisition were selected for analysis. For measurements of DRSH-1::GFP foci, the integrated fluorescence intensity of a region of interest around prominent foci from a single imaging plane was measured using Image J. An adjacent region within the nucleus that lacks DRSH-1::GFP foci was used for background subtraction. To determine the upper limit of focus intensity per nucleus, the first and second brightest foci from each nucleus were identified and quantified. The dimmest foci in the nucleus occasionally yielded negative values after measurement because their intensity was so close to the noise floor. These foci were excluded from the “All foci” analysis. As a result, “All foci” represents all foci that could be identified and measured. Fluorescence intensity was normalized to the average intensity of the comparable (“brightest or “all”) foci in nuclei from wild type DRSH-1::GFP worms during each imaging session.

Measurements of the intensity of GFP in the whole nucleus and estimates of percent DRSH-1::GFP residing in foci both utilized the same images as measurements of foci. However, due to movement of the animals during Z-stack acquisition, only a subset of images could be used for these analyses. Sum projections of the Z-slices containing the nucleus were generated, and the fluorescence intensity of a region of interest containing the nucleus was measured. An adjacent region outside of the nucleus, but within the same cell was used for background subtraction. Fluorescence intensity was normalized to the average intensity of the nuclear GFP signal from wild type DRSH-1::GFP worms for each imaging session. To determine the percentage of DRSH-1::GFP nuclear signal that was in the foci, sum projections were made of each focus in a nucleus, and the fluorescence intensity of a region of interest containing each focus in the resulting images were measured. The sum of all the foci in each nucleus was then calculated and the divided by the total nuclear fluorescence determined above.

### Brood size assay and bursting phenotype scoring

Brood size assay was performed as previously described (Kotagama et al., 2024). In brief, L4 animals of each genotype were singled onto individual plates and transferred to fresh plates every 24 h until embryos were no longer produced. Bursting phenotype was assessed within 48 h after animals reached L3 stage.

### RNA extraction

Bulk samples of L4 and adult stage animals were pelleted from 10cm NMG plates using M9 and snap-frozen in 500µl TRIzol (Life Technologies) until use. Adult animals for Figures 4 and 5 were handpicked in the quantity of one hundred per sample into 20 µl M9 buffer in Eppendorf tubes, and 500µl TRIzol was added to each tube before snap freezing. Upon thawing, samples were vortexed at room temperature for 15 min. 85 µl chloroform was added to each sample and mixed thoroughly. Samples were then centrifuged at 16,000×*g* for 10 min at 4°C. The supernatant was transferred to a fresh tube and an equal volume of 25:24:1 phenol:chloroform:isoamyl alcohol pH=4.5 was added to the sample. Samples were then centrifuged at 16,000×*g* for 15 min at 4°C. The upper aqueous phase was transferred to a fresh tube and an equal volume of isopropanol and 1 µl of GlycoBlue (Thermo Fisher Scientific) was added to the aqueous phase and mixed gently. Samples were frozen at −80°C for 30 min and thawed on ice. RNA precipitation was carried out at 20,000×*g* for 30 min at 4°C. The RNA pellets were then washed twice with 70% ethanol with 5 min of centrifugation at 4°C. Approximately 20 µl of RNase-free water was added to resuspend the RNA.

### qPCR

For miRNA qPCR, reverse transcription reactions were performed using miRNA-specific RT primers with the Taqman microRNA Reverse Transcription (RT) Kit (Thermo Fisher Scientific). For all samples, 1.66 µl of total RNA at 5 ng/µl was used in 5 µl RT reactions. RT reactions were then diluted 1:4 and 1.33 µl was used in a 5 µl qPCR reactions using Taqman miRNA probes and the Taqman Universal Mastermix II with UNG (Thermo Fisher Scientific). Reactions were run in three technical replicates on the Applied Biosystems QuantStudio 6.

Primary miRNA qPCR was performed using KAPA SYBR Fast One-Step qRT-PCR Kit on QuantStudio 6 Flex Real-Time PCR System using primers listed in Table S2. In brief, 1.33 µl of total RNA 5 ng/µl was used in a 5µl qPCR reaction. Reactions were run in three technical replicates for each sample.

### Small RNA sequencing and mapping

Small RNA library preparation was adapted from the NEBNext small RNA library prep kit (New England Biolabs) (Vieux et al., 2021). Briefly, spike-ins were added to 600 ng of total RNA prior to library construction for each sample. Size selection was performed after reverse transcription using 8% urea gel to purify 65-75 nt products. PCR was performed with 15 cycles followed by purification using NEB Monarch PCR purification kit to remove primer dimers. Cloned libraries were sequenced on the Illumina NextSeq 2000 platform.

For bias-minimized small RNA cloning, libraries were constructed as previously described with minor modifications (Mandlbauer et al., 2024). Briefly, 1 µg of total RNA was used as input, and both 5′ and 3′ adaptors were added at a final concentration of 1µM. Spike-ins were introduced across a range of concentrations as previously reported (Sakhawala et al., 2025). Cloned libraries were then sequenced on the Illumina NovaSeq 6000 platform.

Sequence analysis was performed on the NIH high performance computing cluster. Adapter sequences were trimmed using fastp (0.24.0) command fastp -L (S. Chen, 2023). The reference genome was customized by combining *C*. *elegans* genome WS280 with an artificial chromosome containing the synthetic spike-in sequences (Donnelly et al., 2022). Mapping was performed using Bowtie2 using the following settings: --end-to-end --sensitive -q -U (Langmead & Salzberg, 2012). BAM files were sorted and indexed using samtools 1.13 (Danecek et al., 2021). Reads were assigned to miRNAs and piRNAs using htseq 0.13.5 (Putri et al., 2022) with the following settings: --mode union --nonunique fraction -a 0. The htseq analysis on miRNAs was performed using a gff file modified from mirGeneDB (Fromm et al., 2015). The htseq analysis on piRNAs was performed using a gff file from https://www.pirnadb.org/ (Piuco & Galante, 2021). miRNA differential gene expression was analyzed using DESeq2 analysis with default settings (Love et al., 2014) using total piRNA reads or spike-in reads for normalization.

For bias-minimized sequencing data, bam files were collapsed using UMI-tools (Smith et al., 2017) and filtered for reads between 16nt and 30nt.

## Data availability

All primary sequence data have been deposited at NCBI Gene Expression Omnibus under accession number GSE289120. Reviewers may access the data using the following token: abqnacikpdgltyd.

## Acknowledgements

We thank members of the laboratory and Baltimore Worm Club for helpful discussions. We thank Geraldine Seydoux for critical feedback on the manuscript. Multiple strains were obtained from the CGC, which is funded by NIH Office of Research Infrastructure Programs (P40 OD010440). WormBase was used for the design and execution of the study (Sternberg et al., 2024). This research was funded by the NIDDK Intramural Research Program (ZIADK075147) and NHLBI Intramural Research Program (ZIAHL006126).

This research was supported by the Intramural Research Program of the National Institute of Diabetes and Digestive and Kidney Diseases (NIDDK) within the National Institutes of Health (NIH). The contributions of the NIH author(s) were made as part of their official duties as NIH federal employees, are in compliance with agency policy requirements, and are considered Works of the United States Government. However, the findings and conclusions presented in this paper are those of the author(s) and do not necessarily reflect the views of the NIH or the U.S. Department of Health and Human Services.”

## Author Contributions

B.Y. designed and performed experiments, analyzed the data, and wrote and revised the manuscript. B.G. performed experiments, analyzed data, and helped revise the manuscript. N.R. designed experiments and helped revise the paper. R.S. designed and performed experiments. K.M designed experiments and wrote and revised the manuscript.

