## Supplemental Figures for "An intrinsically disordered region of Drosha selectively promotes miRNA biogenesis, independent of tissue-specific Microprocessor condensates"

**List of Supplement Materials**

**Figure S1**. Fluorescent tags do not disrupt the molecular function of Microprocessor.

**Figure S2.** More characterization of the nuclear DRSH-1 condensates.

**Figure S3.** DRSH-1::GFP hemizygous animals display dimmer DRSH-1 foci but no defect in miRNA biogenesis.

**Figure S4.** Quantification of DRSH-1::GFP granule intensity.

**Figure S5.** Characterization of *drsh-1(ΔIDR1)* phenotypes and DESeq2 analysis using spike-ins as the normalization method with both NEBNext and bias-minimized protocols.

**Figure S6.** isomiR distribution analysis comparing ΔIDR2 mutant to wild type.

**Figure S7.** Pri-miRNA features are not predictive of IDR1 dependence.

**Figure S8.** *mir-80* variant loci generate canonical *mir-80*.

**Figure S9.** miRNA-specific effects among clustered miRNAs.

**Table S1.** Strains used in this study.

**Table S2.** Alleles generated and oligonucleotides used in this study.

**Table S3.** Samples used for deep sequencing.

**Table S4.** Raw miRNA and spike-in reads for all samples.

**Table S5.** DESeq2 analysis results of small RNA sequencing comparing *drsh-1(ΔIDR1)* versus wild type at L4 stage using piRNAs reads as normalization method.

**Table S6.** DESeq2 analysis results of small RNA sequencing comparing *drsh-1(ΔIDR1)* versus wild type at adult stage using piRNAs reads as normalization method.

**Table S7.** DESeq2 analysis results of small RNA sequencing comparing *drsh-1(ΔIDR2)* versus wild type at L4 stage using piRNAs reads as normalization method.

**Table S8.** DESeq2 analysis results of small RNA sequencing comparing *drsh-1(ΔIDR2)* versus wild type at adult stage using piRNAs reads as normalization method.

**Table S9.** DESeq2 analysis results of small RNA sequencing comparing *drsh-1(ΔIDR1)* versus wild type at L4 stage using spike-ins as normalization method.

**Table S10.** DESeq2 analysis results of small RNA sequencing comparing *drsh-1(ΔIDR1)* versus wild type at adult stage using spike-ins as normalization method.

**Table S11.** DESeq2 analysis results of small RNA sequencing comparing *drsh-1(ΔIDR2)* versus wild type at L4 stage using spike-ins as normalization method.

**Table S12.** DESeq2 analysis results of small RNA sequencing comparing *drsh-1(ΔIDR2)* versus wild type at adult stage using spike-ins as normalization method.

**Table S13**. DESeq2 analysis results of small RNA sequencing comparing *drsh-1(ΔIDR1)* versus wild type at adult stage using spike-ins as normalization method (bias-minimized small RNA cloning protocol).

**Table S14**. DESeq2 analysis results of small RNA sequencing comparing *drsh-1(ΔIDR2)* versus wild type at adult stage using spike-ins as normalization method (bias-minimized small RNA cloning protocol).

**Table S15.** DESeq2 analysis results of small RNA sequencing comparing *drsh-1::gfp;tdp-1::mScarlet* versus *drsh-1::gfp;tdp-1::mScarlet/drsh-1(null)* at adult stage using piRNAs as normalization method.

**Table S16**. DESeq2 analysis results of small RNA sequencing comparing *drsh-1::gfp* versus wild type at adult stage using spike-ins as normalization method.

**Table S17**. DESeq2 analysis results of small RNA sequencing comparing *pash-1::mScarlet* versus wild type at adult stage using spike-ins as normalization method.

**Table S18**. DESeq2 analysis results of small RNA sequencing comparing *drsh-1::gfp pash-1::mScarlet* versus wild type at adult stage using spike-ins as normalization method.

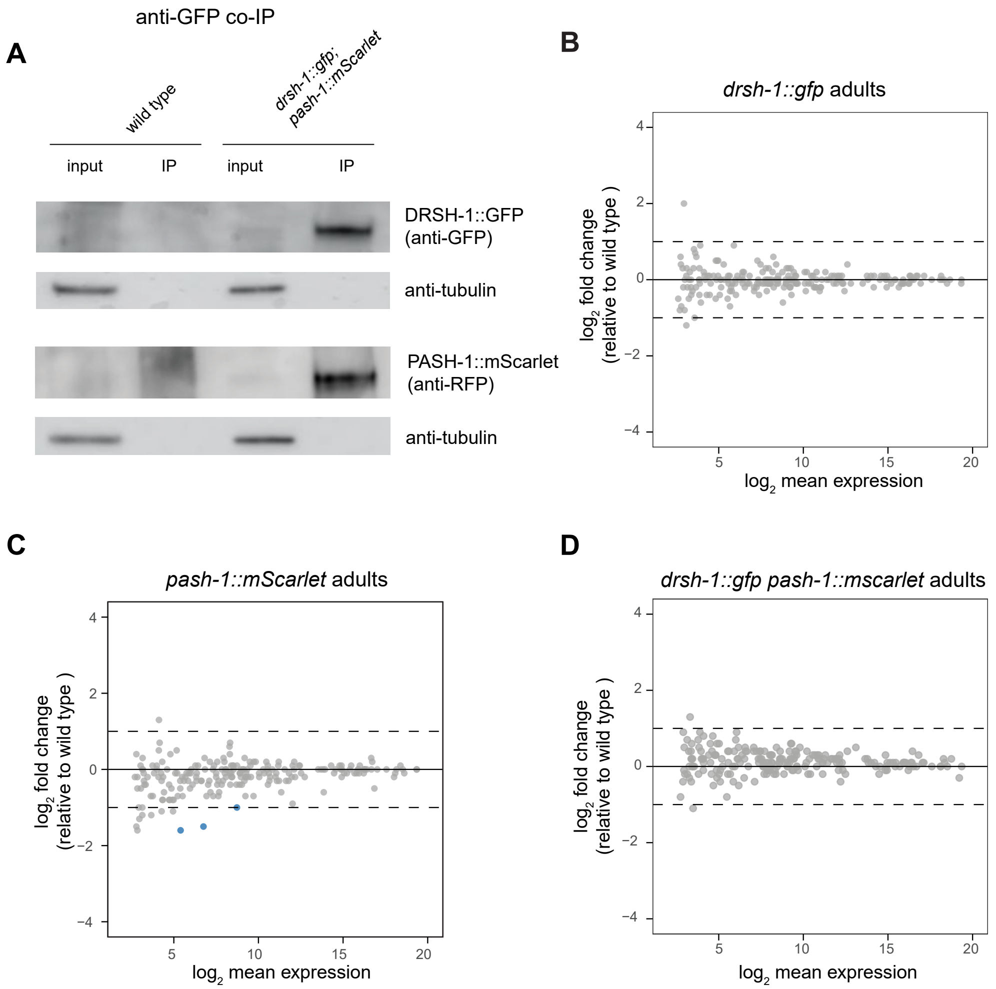


**Figure S1. Fluorescent tags do not disrupt molecular function of Microprocessor.** (A) Immunoprecipitation showing the interaction between DRSH-1::GFP and PASH-1::mScarlet is preserved in the fluorescently tagged strain. 0.5% of total lysate was loaded in each input lane and 20% of IP content was loaded in IP lanes. (B-D) DESeq2 analysis of small RNA sequencing on the different fluorescently tagged strains in comparison to wild type. Spike-ins are used as normalization method.

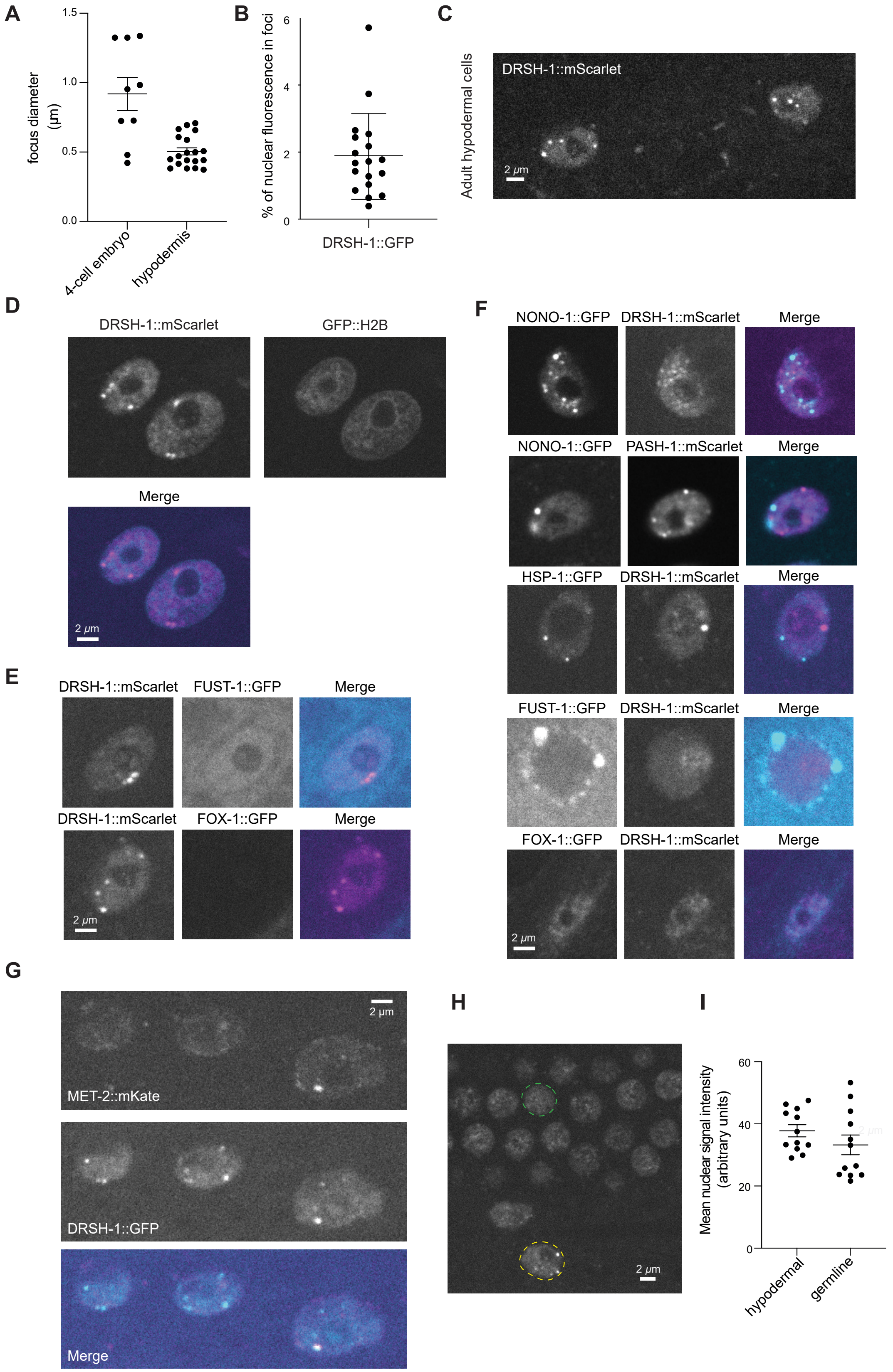


**Figure S2. More characterization of the nuclear DRSH-1 condensates.** (A) Condensate sizes measured in early embryos and hypodermal cells. (B) Quantification of percentage of DRSH-

1::GFP located in granules in hypodermal cells. (C) Representative image of DRSH-1::mScarlet in two hypodermal nuclei. Maximum intensity projection. (D) Co-labeling in hypodermal cells of DRSH-1::mScarlet and GFP::H2B. (E) Co-labeling in hypodermal cells focusing on DRSH-1::mScarlet foci. Single focal plane shown. (F) Co-labeling focusing on foci of the GFP fusion protein. FUST-1::GFP granules are located in germ cells; FOX-1::GFP is visible in muscle cells. Single z plane shown. (G) Representative images of MET-2::mKate imaged together with DRSH-1::GFP in hypodermal cells. Single z plane shown. (H) Representative image showing hypodermis nuclei along with germ cell nuclei in the same focal plane. Yellow dotted line circles a hypodermis nucleus; green dotted line outlines a germ cell nucleus. (I) Quantification of mean nuclear signal comparing hypodermal cells to germ cells.

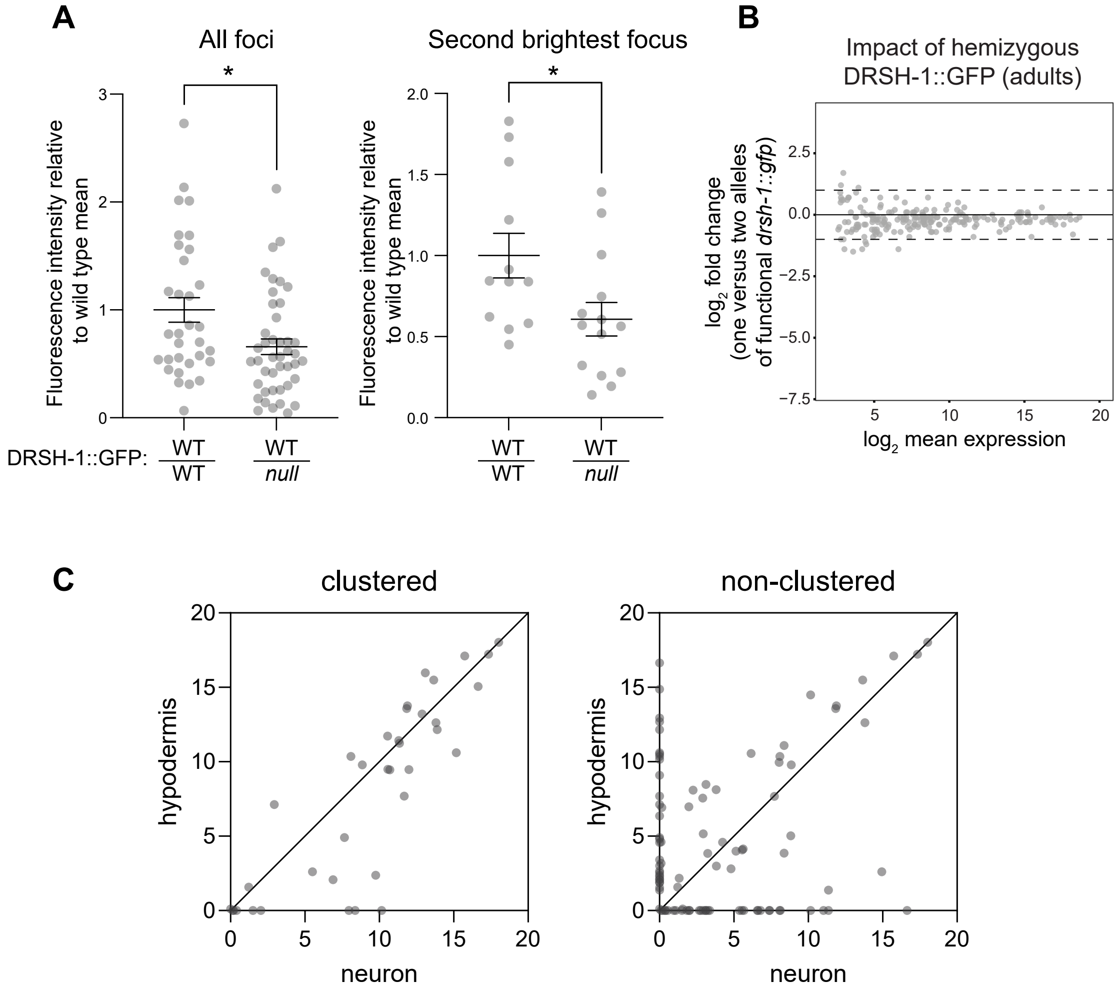


**Figure S3. DRSH-1::GFP hemizygous animals display dimmer DRSH-1 foci but no defect in miRNA biogenesis.** (A) Quantification of the signal intensity of all foci (left) or the second brightest focus per nucleus (right) across indicated genotypes in hypodermal cells. Signal is normalized to the average signal in the control genotype (homozygous DRSH-1::GFP). N ≥ 6 animals per genotype. Two tailed unpaired Student's t-test, * $p<0.05$. (B) MA plot showing the DESeq2 analysis result of adult miRNA sequencing comparing *drsh-1::gfp/drsh-1(null)* to homozygous *drsh-1::gfp* using total piRNA reads as the normalization method. X-axis shows the $\log_2$ average miRNA abundance and Y-axis shows the $\log_2$ fold change of miRNA level. (C) Comparison of miRNA expression levels from clustered or single miRNA loci in condensate forming (hypodermis) or condensate lacking (neuron) tissues. Values are TPM as reported in Wang, et al. *Nat Commun* 2024.

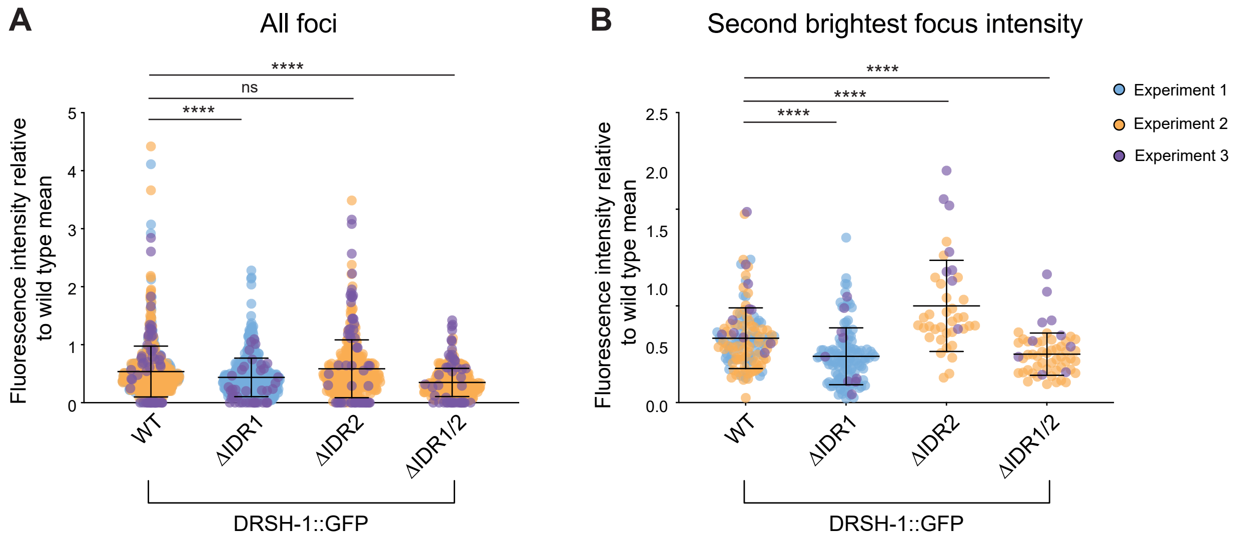


**Figure S4. Quantification of DRSH-1::GFP granule intensity.** Super plots showing the quantification of all foci (A) and second brightest focus per nucleus (B). Signal is normalized to the average signal in the control genotype (wild type DRSH-1::GFP) from the same imaging session. Dots are color-coded according to imaging session. $N \geq 15$ animals per genotype combining three experiments. Two tailed unpaired Student's t-test, * $p<0.05$, ** $p<0.01$, *** $p<0.001$, **** $p<0.0001$.

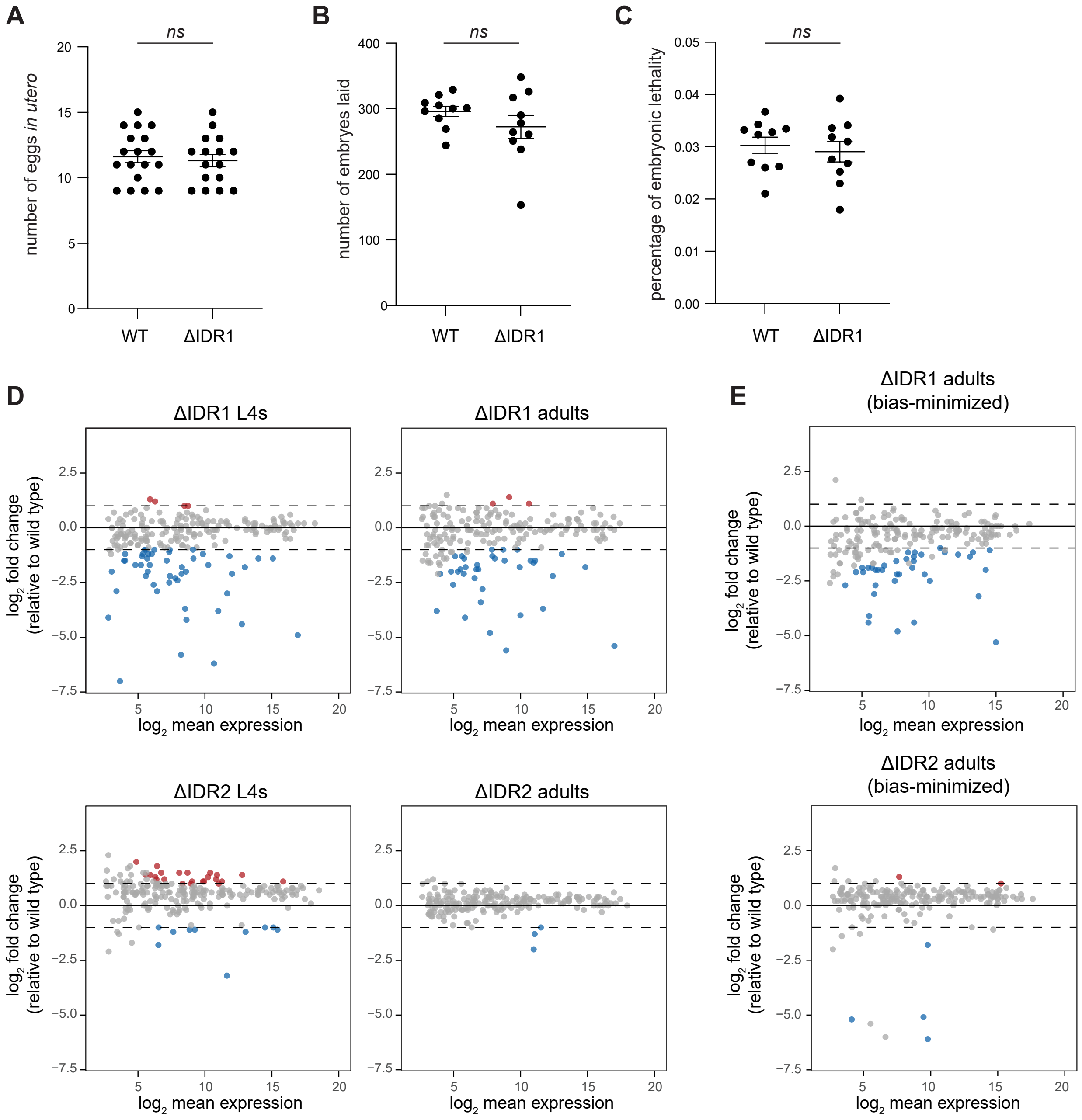


**Figure S5. Characterization of *drsh-1(ΔIDR1)* phenotypes and DESeq2 analysis using spike-ins as the normalization method.** (A) Number of eggs *in utero* in adult hermaphrodite animals 24 hours post L4 synchronization. (B) Brood size assay comparing *drsh-1(ΔIDR1)* and wild type at 20˚C. (C) Embryonic lethality assay comparing *drsh-1(ΔIDR1)* and wt at 20˚C. Two tailed unpaired Student's t-test, ns not significant. (D) DESeq2 analysis using spike-ins as the normalization method. Top two panels show the comparison between ΔIDR1 and wild type, and the bottom two panels show the comparison between ΔIDR2 and wild type. Results are very similar to normalization by total piRNA reads. (E) Bias-minimized small RNA sequencing comparing ΔIDR1 to wild type (top) or ΔIDR2 and wild type (bottom) with spike-in normalization.

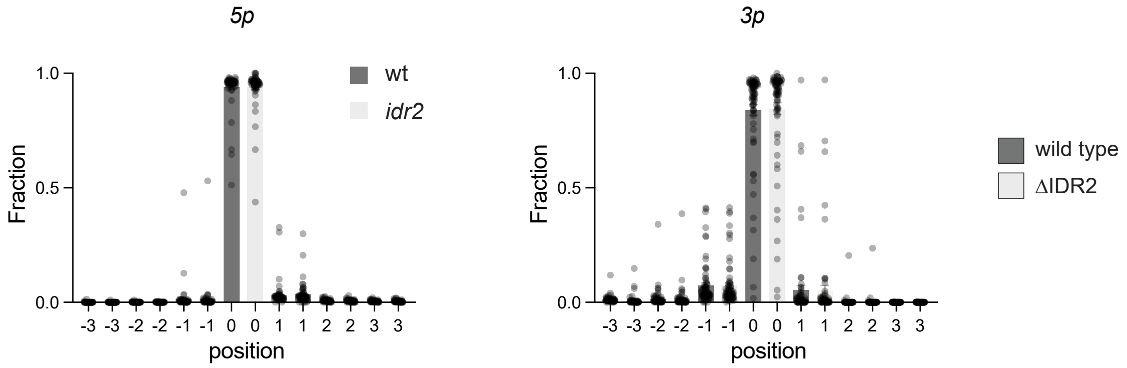


**Figure S6. isomiR distribution analysis comparing ∆IDR2 mutant to wild type.** Distribution of terminal miRNA positions for 5′ ends of 5p miRNA strands and 3′ ends of 3p strands in wild type and *drsh-1(∆IDR2)* animals at adult stage. No significant differences were observed for total or individual miRNAs.

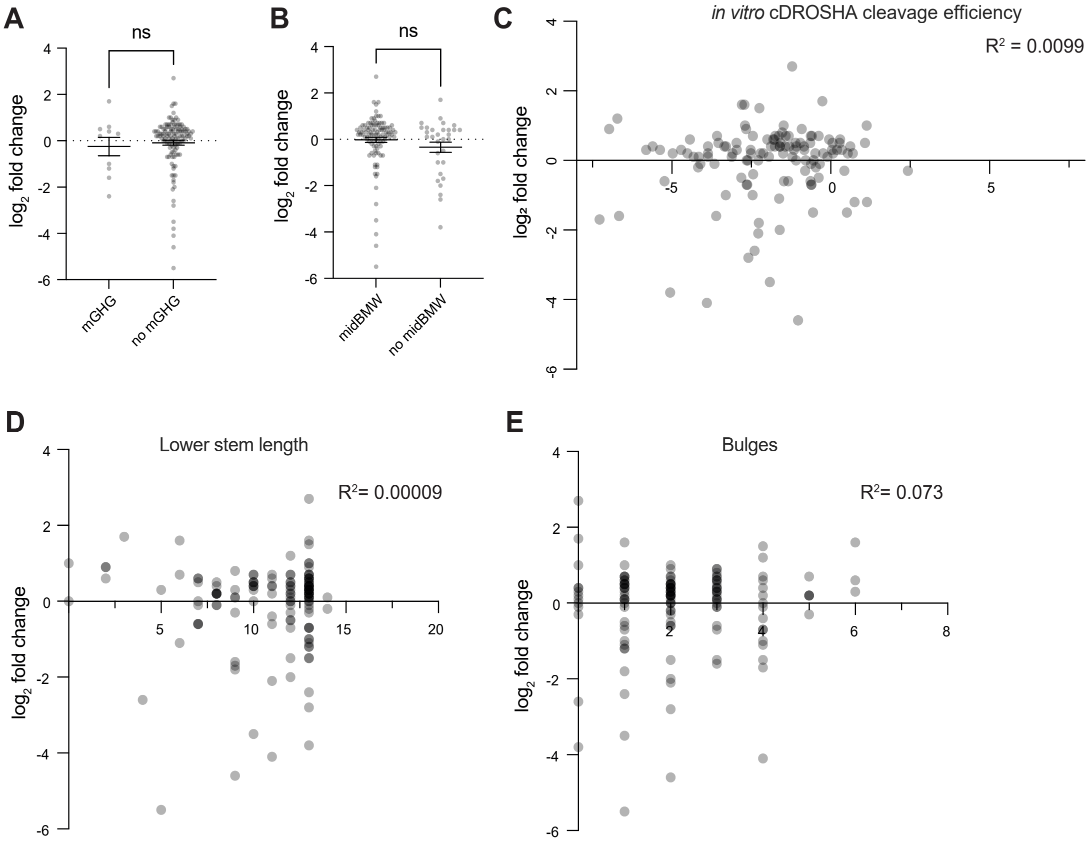


**Figure S7. Pri-miRNA features are not predictive of IDR1 dependence.** (A and B) IDR1 sensitivity between miRNA groups based on classifications: containing or lacking the mGHG motif (A) or the midBMW motif (B). Two tailed unpaired Student's t-test, ns, not significant. (C) X-Y plot showing correlation between IDR1 sensitivity measured by miRNA $log_2$ fold change (ΔIDR1/wild type L4) on the Y-axis and *in vitro C. elegans* Drosha cleavage efficiency from Nguyen, et al. 2023 on the X-axis. (D-E) X-Y plots showing correlation between IDR1 sensitivity measured by miRNA $log_2$ fold change (ΔIDR1/wild type) on the Y-axis and lower stem length (D) or number of bulges (E) on the X-axis.

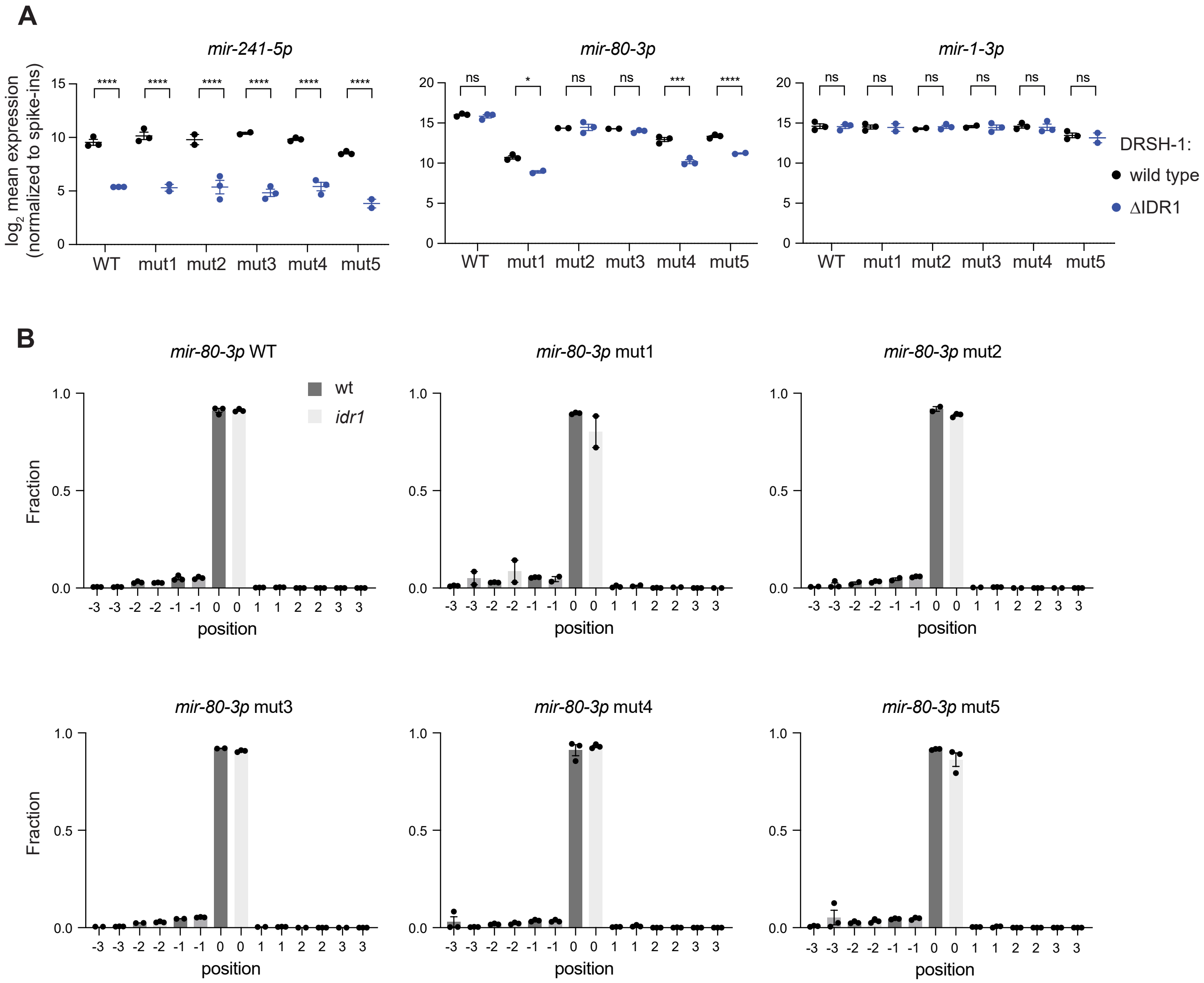


**Figure S8. *mir-80* variant loci generate canonical *mir-80*.** (A) Spike-in normalized miRNA abundance of *mir-80* recapitulates miRNA Taqman qPCR results. *mir-1* and *mir-241* retain IDR1-independence or -dependence, respectively, in all *mir-80* variant backgrounds. (B) IsomiR distribution analysis of the *mir-80*-3p 3' end position in different *mir-80* variant backgrounds.

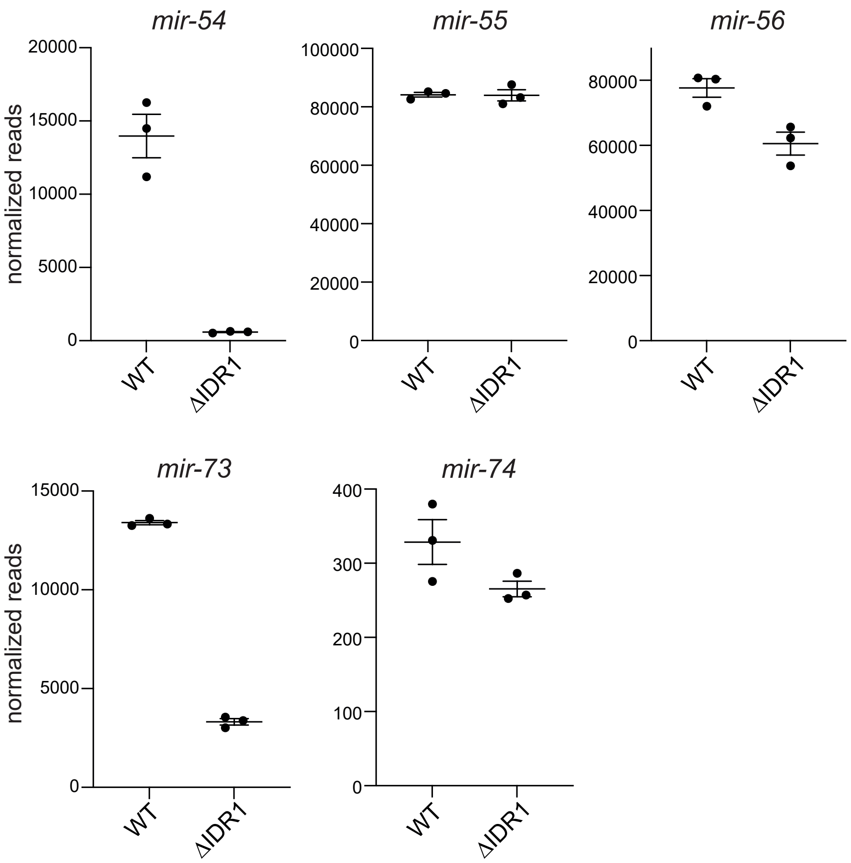


**Figure S9. miRNA-specific effects among clustered miRNAs.** miRNA abundance changes in the *mir-54/55/56* and *mir-73/74* clusters in L4 samples comparing *drsh-1(wild type)* and *drsh-1(*Δ*idr1)*. Data points were extracted from DESeq2 analysis results that were shown in Figure 4.
